# P-glycoprotein exofection between fetal and maternal cells as a mechanism of intercellular material transfer at the feto maternal interface

**DOI:** 10.64898/2026.01.04.697556

**Authors:** Madhuri Tatiparthy, Amanda Wang, Vineeth Mahajan, Pilar Flores-Espinosa, Emmanuel Amabebe, Tilu Jain Thomas, Xiao-Ming Wang, Lauren S Richardson, Ramkumar Menon, Ananth K Kammala

## Abstract

Cells can recover lost protein functions through a process we term exofection, in which extracellular vesicles deliver functional molecular cargo to recipient cells and transiently reprogram their activity. Here we show that exosomes derived from fetal chorion trophoblast cells (CTCs) restore P-glycoprotein (P-gp) efflux transporter function in inflammation-impaired maternal decidual cells (DECs) at the feto-maternal interface. CTCs maintain high P-gp expression under inflammatory stress, whereas DECs exhibit marked downregulation of transporter genes and proteins. Proteomic analysis revealed that CTC-derived exosomes package P-gp as a stable cargo that enters DECs through clathrin- and heparan sulfate–dependent uptake pathways. Delivery of CTC exosomes reinstated P-gp abundance and efflux capacity in LPS-stimulated or P-gp-deficient DECs, as shown by calcein efflux and immunofluorescence assays. In pregnant P-gp knockout mice, exosome treatment restored systemic clearance of the P-gp substrate tacrolimus and improved pharmacokinetic parameters. These findings establish exofection as a naturally occurring mechanism of transporter rescue at the feto-maternal interface, where fetal exosomes compensate for inflammation-induced maternal loss of efflux capacity. By restoring P–gp–mediated barrier function, exofection provides a protective strategy that limits the accumulation of xenobiotics and cytokines in maternal tissues and safeguards the fetus. This work reveals a previously unrecognized form of intercellular communication with broad implications for fetal protection, placental biology, cellular engineering, and the delivery of therapeutic proteins.

**One Sentence Summary:** Exofection is identified as a novel mechanism of transient cellular engineering via exosome-mediated functional protein delivery between cells.

## INTRODUCTION

Exosomes are essential mediators of intercellular communication during human pregnancy, coordinating signaling between fetal and maternal tissues to maintain homeostasis throughout gestation and parturition. Beyond their classical roles in cargo transport, we recently identified a specialized form of exosome-mediated functional transfer called “exofection(*1*),” which is defined as the delivery of a functional protein cargo to a recipient cell that has lost the ability to express or utilize that protein(*2, 3*). Through exofection, recipient cells regain transient, non-genetic restoration of protein function, allowing tissues to adapt to dynamic physiological demands(*3, 4*). This endogenous mechanism has been proposed to operate across distinct physiologic compartments, particularly between the fetus and mother, where localized disruptions in cellular function must be rapidly corrected to preserve pregnancy.

The feto-maternal interface (FMi) represents a critical anatomical and immunological environment where maternal and fetal tissues cooperate to maintain homeostasis for successful pregnancy(*5, 6*). Two interfaces exist: the well-characterized placenta-decidua basalis and the broader fetal membrane–decidua parietalis (choriodecidua), the latter formed by fetal chorionic trophoblast cells (CTCs) in comparison to maternal decidual stromal cells (DECs). This choriodecidual interface spans a large surface area and plays essential roles in immune regulation, cytokine buffering, tissue remodeling, and mechanical containment(*6–11, 12, 13*). Disruption of these processes, primarily through inflammatory activation, contributes to adverse pregnancy outcomes, including recurrent miscarriage, preeclampsia, and spontaneous preterm birth(*14, 15*). Maintaining molecular and functional homeostasis at the FMi is therefore fundamental to fetal protection(*5, 12, 16*).

Transporter proteins (TPs), such as P-glycoprotein (P-gp/ABCB1) and breast cancer resistance protein (BCRP/ABCG2), are essential components of the defense system at the FMi. These ATP-binding cassette transporters regulate the efflux of xenobiotics, endogenous metabolites, inflammatory mediators, and oxidative byproducts, functioning as molecular gatekeepers at epithelial and endothelial barriers throughout the body (*18–20*). During pregnancy, TPs are highly concentrated on both fetal and maternal sides of the FMi (*21–23*), where they prevent harmful substances from entering fetal circulation and help reduce cytokine-driven tissue damage (*24, 25*). Among these, P-gp plays a key role in preserving epithelial integrity(*24, 25*), limiting drug buildup(*26, 27*), and maintaining the barrier function necessary for normal gestation. However, transporter expression is flexible and highly responsive to environmental signals; physiological and pathological inflammation can significantly impact P-gp levels and activity, with important implications for maternal and fetal health (*18, 19, 28, 29*).

A growing body of evidence suggests that inflammation can downregulate P-gp in multiple barrier tissues; however, its regulation at the FMi remains unclear. Our findings demonstrate that this effect is highly cell-type specific, occurring in maternal decidual stromal cells (DECs) but not in fetal chorionic trophoblast cells (CTCs), revealing a previously unrecognized functional asymmetry at the interface. In DECs, the engagement of TLR4 by endotoxin or endogenous danger signals activates both canonical and non-canonical NF-κB pathways, inducing a cascade of proinflammatory cytokines, including IL-1β, TNF, IL-6, GM-CSF, and interferon-stimulated genes. NF-κB activation is known to interfere with nuclear receptors such as PXR and CAR, which generally support ABCB1 transcription, leading to rapid suppression of P-gp expression. In parallel, inflammatory nitric oxide and oxidative stress promote ROS-dependent internalization and proteasomal degradation of P-gp, further reducing efflux capacity. Our transcriptomic analyses of primary FMi cells support this mechanistic framework, revealing the robust activation of NF-κB-, IL-17-, and interferon-associated pathways specifically in DECs, accompanied by significant downregulation of ABCB1. In contrast, CTCs exhibit attenuated inflammatory responsiveness, preserve ABCB1 transcription, and maintain P-gp function even under identical inflammatory stimuli. This differential regulation of P-gp in DECs and CTCs is a novel observation from our laboratory, and to our knowledge represents the first evidence of inflammation-induced transporter asymmetry across the human FMi.

This asymmetric vulnerability has significant physiological implications. Loss of P-gp–mediated efflux on the maternal side of the FMi increases tissue permeability to xenobiotics, inflammatory mediators, and metabolic toxins, potentially intensifying local inflammation and weakening fetal protection. However, most inflammatory exposures during pregnancy, such as transient microbial encounters, redox fluctuations, or cytokine waves, do not result in adverse outcomes or preterm birth. This discrepancy suggests that the body has endogenous compensatory mechanisms that maintain efflux balance when maternal transporter levels drop. We hypothesize that the fetus actively participates in this compensation by releasing CTC-derived exosomes, which are rich in functional P-gp and are then internalized by maternal DECs. Through exofection, these exosomes deliver P-gp protein to maternal cells, restoring efflux capacity and strengthening barrier function during inflammatory stress.

While multiple ABC transporters contribute to efflux at the FMi, we selected P-gp for in-depth investigation because it represents the primary efflux barrier for xenobiotics, cytokine metabolites, and oxidative byproducts in both fetal and maternal tissues. Importantly, P-gp exhibits the most significan**t** dynamic regulation in response to inflammation, undergoing rapid transcriptional suppression and protein degradation in maternal DECs, whereas it remains stable in CTCs. This distinct regulatory behavior provides a clear mechanistic framework to evaluate whether exosome-mediated protein transfer can functionally rescue a transporter that is both essential and inflammation-sensitive. In this study, we test the hypothesis that chorion-derived exosomes can restore P-gp expression and function in maternal DECs subjected to inflammation in vitro and extend this mechanism to an in vivo physiological context using pregnant P-gp knockout mice. Our findings support a model in which exofection serves as a biologically encoded, transient repair mechanism, enabling the fetus to compensate for maternal loss of transporter function and sustain homeostasis at the FMi.

## RESULTS

### Differential expression and directional efflux function of P-gp at the FMi

The anatomical structure of the fetal membrane–decidual interface (FMi) is illustrated in **Figure 1A**, where the amnion epithelial cell (AEC) layer covers the chorionic trophoblast cell (CTC) layer, which directly contacts the decidua parietalis. We first evaluated whether this interface supports functional P-glycoprotein (P-gp) efflux activity using a bidirectional tacrolimus transport **assay (Figure 1B,D).** Tacrolimus, a well-known P-gp substrate (*30*), was added to either the fetal-facing (AEC apical) or maternal-facing (DEC apical) chamber to assess directional transport.

**Figure 1:**
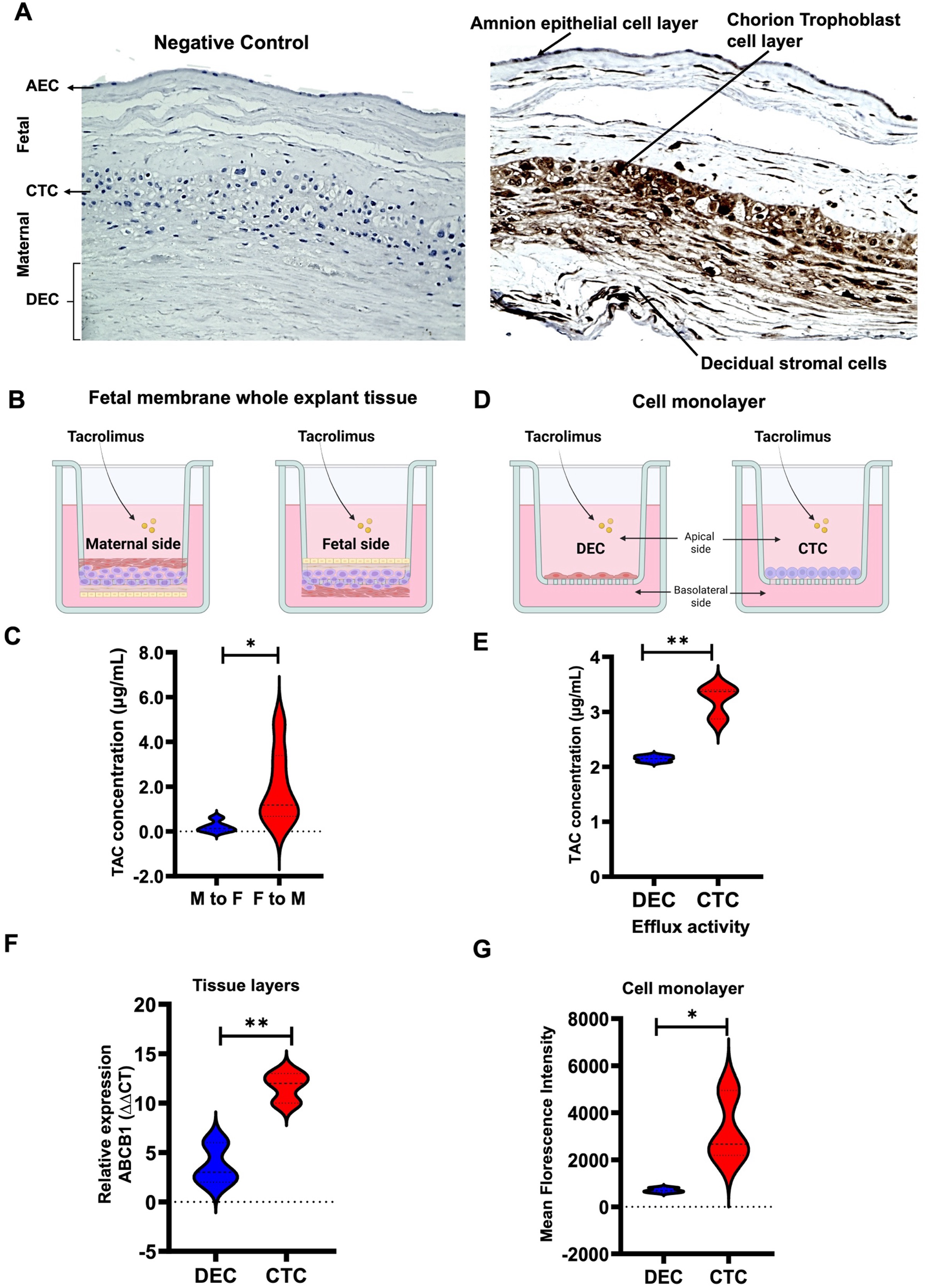
P-gp expression and efflux function at the feto-maternal interface. (A) Immunohistochemical localization of P-gp in human fetal membrane tissues, showing the amnion epithelial layer (AEC), underlying chorionic trophoblast cells (CTCs), and maternal decidua. Representative control (left) and P-gp–stained sections (right) are shown at 10× magnification. (B) Schematic of the transwell configuration used for fetal membrane explant studies, with the fetal side oriented apically (left) or the maternal decidual side oriented apically (right). (C) Tacrolimus transport across fetal membrane explants quantified by LC–MS/MS, demonstrating directional efflux from the fetal to the maternal compartment. (D) Transwell systems using isolated CTCs (left) and DECs (right) to assess cell-specific transporter activity. (E) Tacrolimus efflux measured in CTC and DEC monolayers, showing higher efflux capacity in CTCs. (F) Relative ABCB1 mRNA expression in CTCs and DECs determined by qPCR. (G) Flow cytometric quantification of intracellular P-gp protein levels in DECs and CTCs using PE-conjugated antibody staining. Data are presented as mean ± SEM. Statistical analysis was performed using Student’s t-test. *p* < 0.05, **p** < 0.01, \***p** < 0.001.

At early time points (≤2 h), drug movement across the tissue was minimal in both directions. However, by 3 h, tissues mounted with the fetal side facing apically showed a significant increase in tacrolimus transport (p < 0.01), while the maternal-to-fetal orientation displayed negligible transfer **(Figure 1C).** These results reveal a strong directional preference for fetal-to-maternal efflux, consistent with a protective barrier that limits maternal exposure of the fetus to xenobiotics.

To determine whether this directional behavior reflects inherent differences between fetal and maternal cells, we next examined isolated CTC and DEC monolayers. Both cell types demonstrated P-gp–mediated transport, but CTCs exhibited significantly higher efflux capacity (p < 0.01), suggesting that fetal trophoblasts play a larger role in efflux function at the FMi (**Figure 1E).**

We then examined P-gp expression at the protein and transcript levels. Immunohistochemistry revealed strong P-gp staining in the CTC layer, with substantially lower expression in AECs and DECs. Consistent with this pattern, ABCB1 mRNA levels were significantly higher in isolated CTCs compared to DECs **(p < 0.05; Figure 1F).** Flow cytometric analysis further validated this finding, showing significantly higher surface P-gp levels in CTCs than in DECs (p < 0.05; **Figure 1G).** The full gating strategy for flow cytometry analyses is shown in ***Supplementary Figures 1–2***. Overall, these results demonstrate that the fetal side of the FMi has higher P-gp expression and significantly greater efflux activity than the maternal side, creating a physiologically relevant asymmetry in transporter function.

### Transcriptomic profiling reveals inflammation-induced suppression of ABC transporters in DECs and preservation in CTCs

To investigate the changes to the TPs at the FMi during an inflammatory environment, both DECs and CTCs were treated with LPS and subjected to transcriptomic analysis, followed by a thorough characterization of changes in the expression of inflammatory and transporter proteins in each cell type. LPS induced significant activation of inflammatory signaling pathways in DECs following LPS treatment. Notably, Trim21, interferon alpha/beta signaling, and IL-1 signaling pathways were upregulated (**Figure 2A**). In contrast, CTCs displayed minimal transcriptional changes in response to LPS, with no significant activation of canonical inflammatory pathways. Differential gene expression analysis supported these findings, showing substantial upregulation of key inflammatory genes in DECs, including *MMP3, IL6, CXCL1, CXCL8, IFI6, ISG15*, and *HERC6*. These genes are well-known mediators of inflammation and immune cell recruitment. Conversely, in CTCs, while some genes such as *KRT7, UPP1, TAGLN, CXCL8*, and *SP6* were detected, their expression changes were not statistically different (**Figure 2B**). This data supports our earlier reports showing that CTCs are refractory to exogenous stimuli and provide a formidable FMi barrier(*31*).

**Figure 2:**
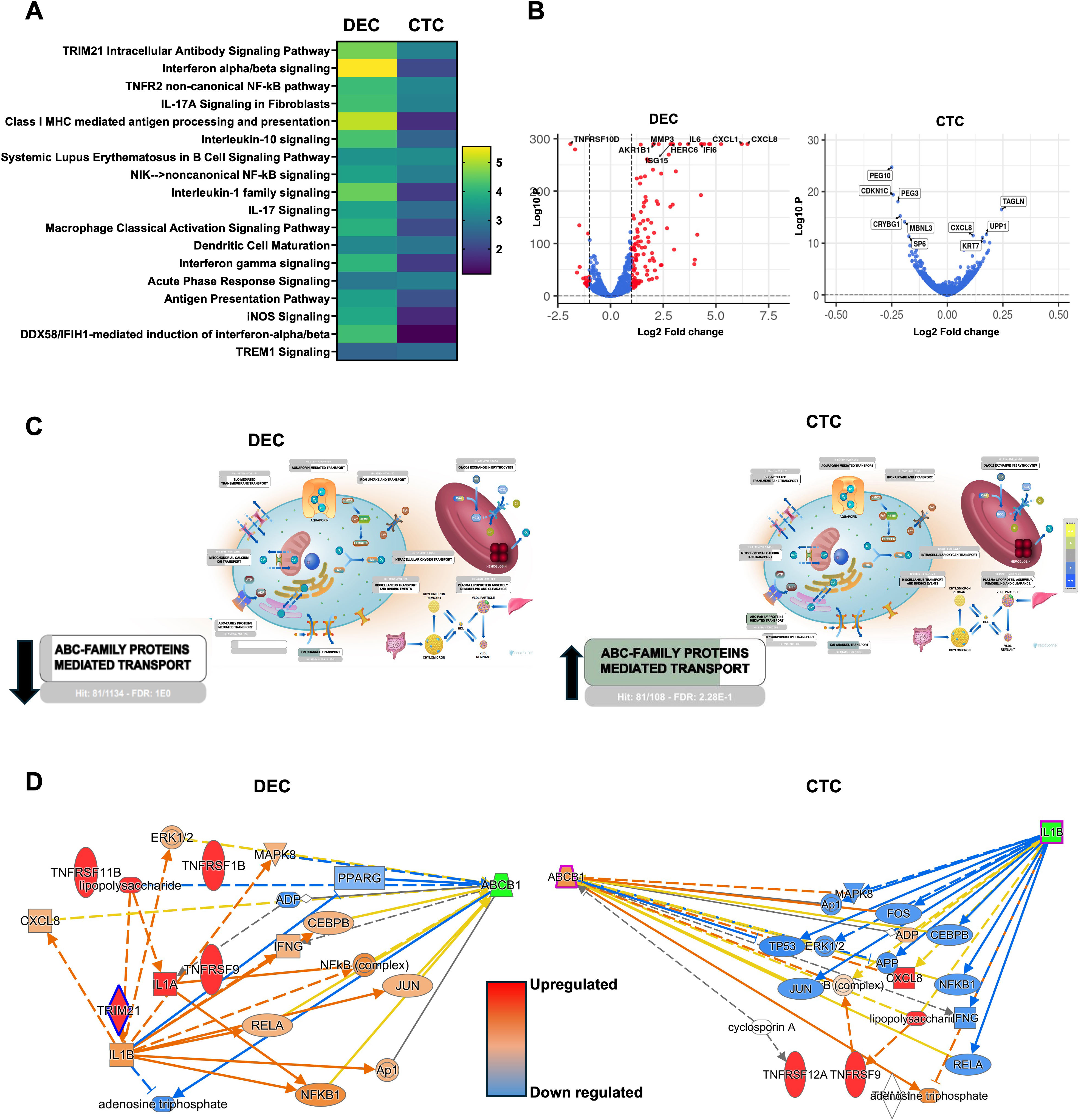
Differential inflammatory responses and transporter regulation in DECs and CTCs following LPS stimulation. (A) Pathway enrichment analysis of differentially expressed genes (DEGs) in LPS-treated DECs (left) and CTCs (right), showing robust activation of immune and inflammatory signaling pathways in DECs and minimal pathway induction in CTCs. (B) Volcano plots depicting significantly upregulated and downregulated genes in DECs (left) and CTCs (right). Key inflammatory mediators are highlighted. (C) Reactome analysis identifying transporter-related pathways altered by inflammation, including ABC transporter families, with ABCB1 downregulated in DECs and preserved or upregulated in CTCs. (D) Network analysis of upstream regulators and signaling molecules associated with ABCB1 expression under inflammatory conditions in DECs (left) and CTCs (right).

To evaluate how this inflammatory asymmetry affects transporter biology, we examined Reactome pathways enriched in DEC versus CTC transcriptomes. The analysis revealed distinct patterns of transporter protein expression between the two cell types. Both DECs and CTCs expressed various transporters, including SLC family members, aquaporins, mitochondrial iron transporters, and ion channels. Of particular interest, the ABC transporter family, including *ABCB1*, was found to be significantly upregulated in CTCs. At the same time, *ABCB1* expression was downregulated in DECs under inflammatory conditions (**Figure 2C)**, left is DEC, and right is CTC. To explore the mechanisms behind *ABCB1* downregulation in DECs, Ingenuity pathway analysis (IPA), upstream regulatory network analysis, and subcellular localization of gene networks were performed (**Figure 2D, *supplementary figures 3 & 4***). This analysis identified several upregulated inflammatory mediators that may contribute to *ABCB1* suppression in DECs, including *TNFRSF11B, IL1B, TNFRSF9, IL1A, TRIM21, ERK, MAPK8, RELA, JUN*, and *NFKB1*. These factors are known to modulate transcriptional activity and pro-inflammatory signaling, suggesting a coordinated suppression of *ABCB1* in inflamed DECs.

Conversely, in CTCs, inflammatory markers such as TNFRSF12A, TNFRSF9, and CXCL8 were upregulated, and *IL-1β* expression was reduced after LPS treatment. The downregulation of *IL-1β* may contribute to a muted downstream inflammatory response, potentially preserving *ABCB*1 expression in CTCs under similar LPS exposure. Our transcriptomic and in vitro experimental data determined that LPS stimulation significantly downregulates P-gp/ABCB1 expression in DECs, whereas CTCs remain unaffected. This differential response supports tissue-specific regulation of ABC transporters under inflammatory conditions. In DECs, LPS triggers activation of non-canonical NF-κB signaling pathways, including upregulation of TNF receptor family genes *IL1B, TRIM21*, and *JUN*. These inflammatory responses elevate pro-inflammatory cytokines, such as IL-6, IL-8, and GM-CSF, which are known to suppress *ABCB1* expression and consequently diminish efflux activity (*32–35*). Together, these transcriptomic analyses demonstrate that inflammation induces a strong, cell-type–specific suppression of P-gp/ABCB1 in maternal DECs while sparing fetal CTCs. This asymmetric response provides a mechanistic basis for the differential efflux capacity observed between the two cell types. It supports the hypothesis that exofection may compensate for inflammation-driven loss of transporter function on the maternal side of the interface.

### Inflammatory conditions impair P-gp expression and efflux function in DECs but not in CTCs

To validate the transcriptomic signatures identified in DECs and CTCs (Figure 2), we measured cytokine secretion after LPS stimulation. DECs showed a strong inflammatory response, with significant increases in several cytokines including IL-6, IL-8, GM-CSF, and IL-1β, confirming activation of the canonical NF-κB and interferon-related pathways. In contrast, CTCs showed no measurable change in cytokine production, which is consistent with their muted transcriptomic response and supports the idea that trophoblasts are naturally resistant to inflammatory stimuli at the FMi **(Figure 3A).**

**Figure 3:**
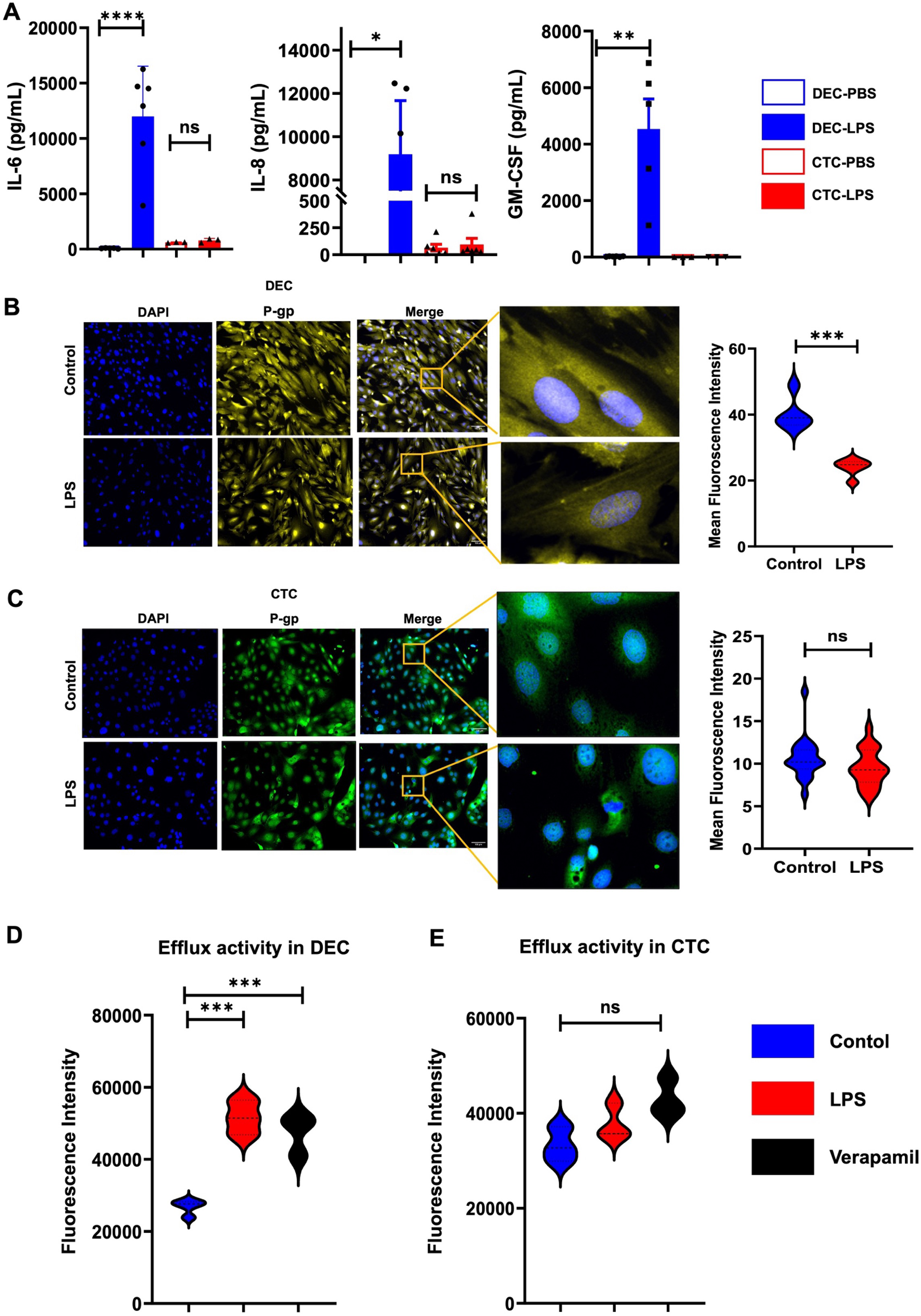
Inflammation selectively suppresses P-gp expression and efflux function in DECs but not CTCs. (A) Multiplex cytokine profiling of IL-6, IL-8, GM-CSF, and TNF in DECs and CTCs following LPS exposure, showing robust inflammatory cytokine release in DECs and minimal induction in CTCs. (B–C) Representative immunofluorescence images of P-gp (cyan) and DAPI (blue) in DECs (B) and CTCs (C) under control and LPS-treated conditions, with corresponding merged images. Bar graphs show quantification of P-gp mean fluorescence intensity. (D–E) Calcein-AM dye accumulation in DECs (D) and CTCs (E) under control, LPS, and verapamil-treated conditions, assessing P-gp–dependent efflux activity. Increased fluorescence denotes reduced efflux. Data are presented as mean ± SEM. Statistical analyses were performed using Student’s t-test or one-way ANOVA. *p* < 0.05, **p** < 0.01, \***p** < 0.001.

We evaluated the influence of inflammation on P-gp protein levels. Immunocytochemistry showed a significant decrease in P-gp signal intensity in DECs after LPS exposure (p < 0.01), evidenced by lower membrane-bound fluorescence and less punctate staining, both indicating reduced transporter density **(Figure 3B).** This decrease aligns with the transcriptional repression and pathway activation observed in DECs, where NF–κB–dependent signaling, TRIM21 induction, and IL-1 family activation lead to the suppression of ABCB1. In contrast, P-gp levels in CTCs remained stable, maintaining both membrane localization and fluorescence intensity similar to untreated controls **(Figure 3C).** These results confirm at the protein level that inflammatory signals selectively impair maternal but not fetal transporter expression.

To assess whether decreased P-gp expression leads to reduced efflux capacity, we conducted calcein-AM accumulation assays. DECs exposed to LPS showed significantly increased intracellular calcein fluorescence (p < 0.01), indicating decreased P-gp–mediated efflux **(Figure 3D). Notab**ly, verapamil treatment further elevated calcein retention under both basal and inflammatory conditions, confirming that the functional decline is P-gp–dependent. The level of calcein accumulation in inflamed DECs was nearly the same as in verapamil-inhibited controls, demonstrating a substantial loss of transporter function.

In CTCs, calcein efflux remained intact despite LPS exposure, with no significant change in fluorescence intensity compared to controls **(Figure 3E).** This preservation of efflux ability matches stable ABCB1 transcription and protein levels in CTCs under inflammatory stress. Overall, these results show that inflammation specifically reduces both P-gp expression and efflux activity in maternal DECs while not affecting fetal CTCs. This functional difference supports our transcriptomic data and emphasizes a significant biological variation in transporter resilience at the FMi, offering a mechanistic explanation for why exofection might act as a backup rescue pathway during inflammatory stress.

### Exofection by fetal exosomes as a mechanism to restore P-gp function on the maternal side

To determine whether fetal trophoblast–derived exosomes contribute to intercellular transfer of transporter proteins at the FMi, we first isolated EVs from CTCs and DECs using a scalable tangential-flow filtration (TFF) platform **(Figure 4A)**. Proteomic profiling revealed notable cell-type–specific differences in transporter cargo. DEC-derived exosomes contained relatively low levels of TPs, whereas CTC-exosomes showed significant enrichment of ABC and SLC family members, especially under inflammatory conditions **(Figure 4B).** The targeted increase in TP cargo within LPS-stimulated CTC-exosomes reflects the transcriptional stability of ABCB1/P-gp observed in CTCs (Figure 2), suggesting a coordinated mechanism where fetal cells maintain transporter function both inside cells and in secreted EVs.

**Figure 4:**
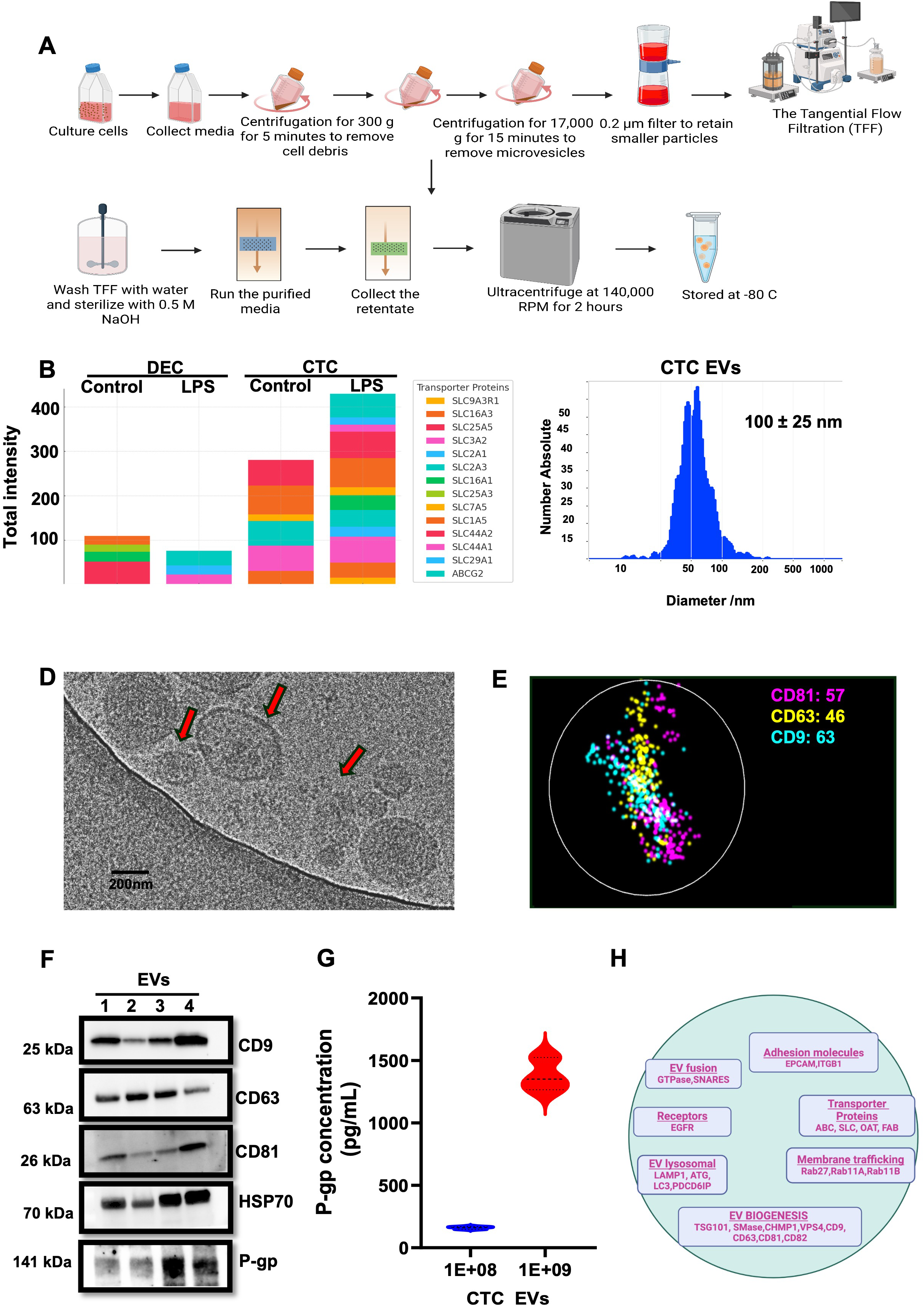
Isolation and molecular characterization of CTC-derived extracellular vesicles (EVs). (A) Schematic overview of the EV isolation and purification workflow from CTC-conditioned media using tangential flow filtration (TFF). (B) Comparative proteomic analysis of transporter proteins in EVs derived from DECs and CTCs under control and LPS-stimulated conditions, highlighting enrichment of transporter families in CTC-EVs. (C) Nanoparticle tracking analysis (Zetaview) showing EV size distribution and particle concentration. (D) Cryo-electron microscopy image demonstrating intact lipid bilayer structures and morphological heterogeneity of EVs. (E) Super-resolution Nanoimager microscopy showing surface expression of canonical EV tetraspanins (CD9, CD63, CD81). (F) Western blot analysis confirming the presence of EV markers (CD9, CD63, CD81), HSP70, purity marker Calnexin, and P-gp cargo in isolated EVs. (G) ELISA-based quantification of P-gp abundance in EV preparations. (H) Functional annotation of proteins identified in CTC-derived EVs, demonstrating enrichment of pathways involved in membrane fusion, vesicle trafficking, receptor-mediated interactions, lysosomal processes, cellular adhesion, and molecular transport.

We then characterized exosomes isolated from unstimulated CTCs to evaluate their structural integrity and cargo-loading capacity. Nanoparticle tracking analysis revealed a mean vesicle diameter of approximately 125 ± 25 nm and particle counts ranging from 80 to 150 particles per frame, indicating consistent and robust EV production **(Figure 4C).** Cryo-EM imaging confirmed a morphologically diverse but structurally intact EV population, including vesicles with single, double, or granular membranes **(Figure 4D; *Supplementary Figure 5*).** These varied morphologies are typical of functional exosomes, which are capable of cargo transport and membrane fusion. To validate exosome identity, we employed single-molecule nanoimaging and immunoblotting of canonical tetraspanins. Nanoimager analysis revealed distinct EV subpopulations expressing CD63, CD9, and CD81 either individually or in dual/triple combinations **(Figure 4E–F).** Western blotting confirmed the presence of these markers and demonstrated the expected enrichment of HSP70 and the absence of the negative control marker Calnexin, indicating minimal contamination from cellular debris.

Importantly, P-gp was consistently detected across multiple exosome batches, and ELISA quantification revealed a positive correlation between vesicle number and P-gp abundance **(Figure 4F–G).** These findings strongly support active, rather than random, loading of P-gp into CTC-derived exosomes.

To understand the molecular basis of this selective cargo enrichment, we analyzed the transcriptome of CTCs during exosome biogenesis. Pathway analysis showed high expression of genes involved in vesicle formation, trafficking, docking, and fusion, including *RAB11A/B, RAB27A, TSG101, VPS4, ESCRT* components, *SNARE* family members, and tetraspanins (*CD63, CD81, CD9*) indicating a mature and transcriptionally enhanced exosome production system **(Figure 4H; *Supplementary Figure 7*).**

Furthermore, CTC-exosomes contained transcripts related to small-molecule transport, adhesion, and endocytosis, along with transporter genes from the ABC, SLC, OAT, and FABP families, highlighting their specialization in intercellular cargo delivery. Together, these data demonstrate that fetal trophoblasts produce exosomes with enhanced transporter content and strong biogenesis signatures, positioning CTC-exosomes as highly effective vesicles capable of transferring functional P-gp to maternal cells. This mechanism, exofection, offers a biologically plausible pathway through which the fetus may compensate for inflammation-induced transporter loss on the maternal side, thereby maintaining fetoprotective efflux capacity at the FMi.

### Maternal DECs efficiently internalize fetal CTC-derived exosomes through defined endocytic pathways

We next examined whether maternal DECs, a prerequisite for P-gp exofection at the FMi, actively internalize fetal CTC-exosomes. PKH-26–labeled exosomes were added to DECs at three physiologically relevant particle concentrations (1×10¹□, 1×10□, 1×10□ particles). Fluorescence imaging showed a clear dose-dependent uptake, with significantly higher intracellular fluorescence at the highest dose (p□<□0.01), indicating robust vesicle internalization **(Figure 5A).**

**Figure 5:**
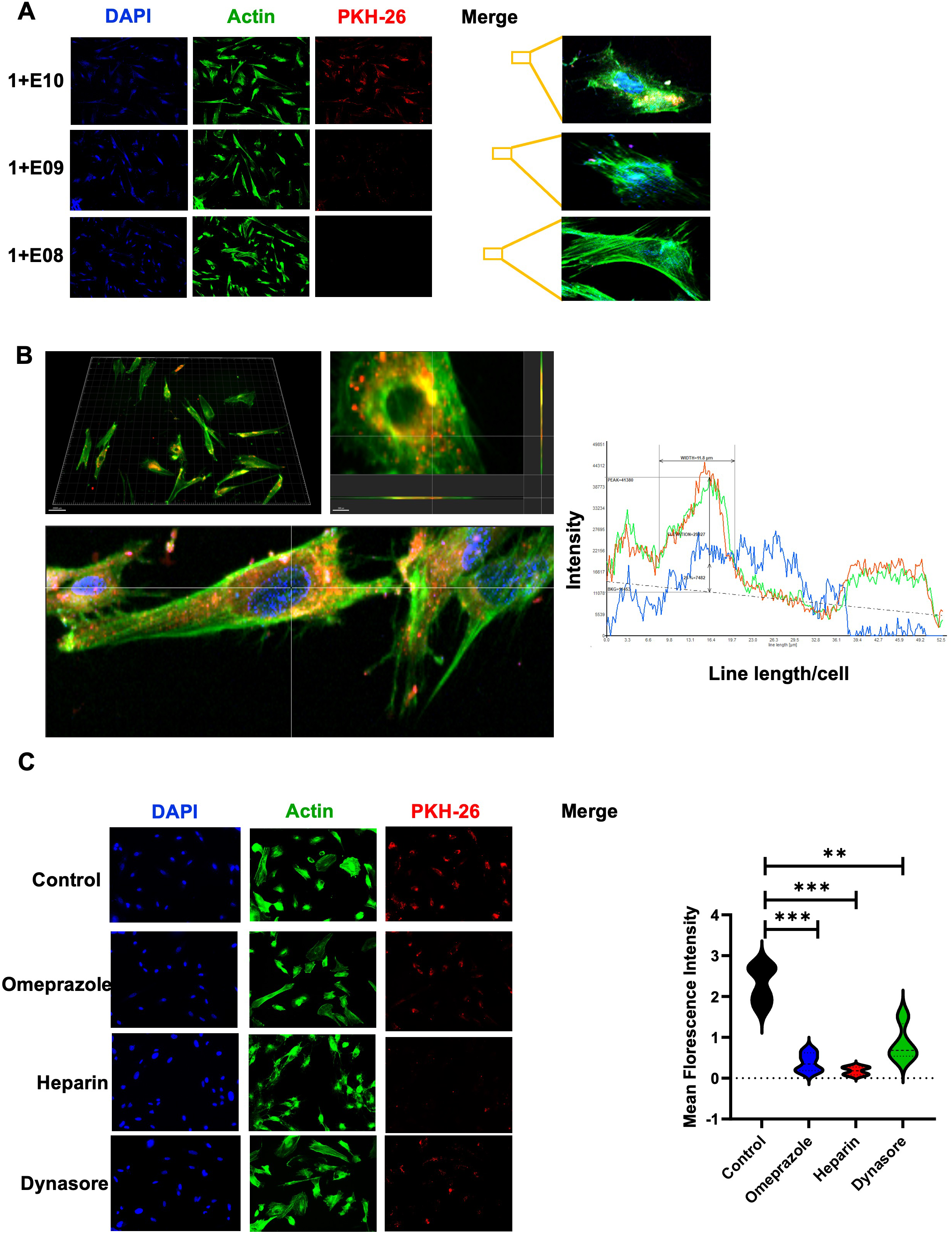
Mechanisms governing maternal DEC uptake of CTC-derived extracellular vesicles. (A) Dose-dependent internalization of EVs by maternal DECs. Cells were treated with increasing EV concentrations (1×10¹□, 1×10□, 1×10□ particles). EVs were labeled with PKH-26 (red), nuclei with DAPI (blue), and actin with phalloidin (green). Quantification of PKH-26 mean fluorescence intensity is shown in the adjacent bar graph. (B) Three-dimensional colocalization of internalized EVs within DECs visualized using z-stack confocal microscopy. Images were reconstructed in Imaris to demonstrate spatial overlap of PKH-26 EV signal with cytoskeletal and nuclear markers, confirming intracellular uptake. (C) Evaluation of uptake pathways using endocytosis-specific inhibitors. DECs were pretreated with pharmacological inhibitors and subsequently incubated with PKH-26–labeled EVs (1×10¹□ particles). Representative fluorescence images and quantitative analysis of PKH-26 intensity demonstrate reduced uptake following inhibition of clathrin-mediated endocytosis, heparan sulfate proteoglycan-mediated binding, and pH-dependent vesicular trafficking. Statistical significance for all assays was determined by Student’s t-test or one-way ANOVA. Data are presented as Mean ± SEM. Significance is indicated as * for p < 0.05, ** for p < 0.01 and *** for p < 0.001

To confirm that the fluorescent signal represented actual intracellular uptake rather than surface attachment, we performed Z-stack imaging on a Keyence system. Orthogonal projections revealed that PKH-26–positive signals were localized within the cytoplasm and colocalized with DAPI and actin structures, confirming the membrane penetration and internalization of CTC-exosomes. Confocal microscopy further supported these findings, displaying precise spatial overlap between exosomal fluorescence and actin filaments, which is consistent with cytoskeletal engagement during vesicle trafficking **(Figure 5B).**

To identify the molecular pathways responsible for exosome entry, we systematically blocked different endocytic routes before exposing the cells to exosomes. DECs were pretreated with inhibitors targeting: dynamin-dependent / clathrin-mediated endocytosis (Dynasore, Chlorpromazine), heparan sulfate proteoglycan–mediated uptake (Heparin), acidification-dependent vesicle maturation and trafficking (Omeprazole), and macropinocytosis/phagocytosis (Amiloride, Wortmannin).

Cytotoxicity assays confirmed that all inhibitor concentrations were non-toxic (Supplementary Figure 8A). Inhibition studies revealed a clear pattern: Dynasore, Heparin, and Omeprazole each caused a significant decrease in exosome uptake (p□<□0.01) **(Figure 5C),** indicating that clathrin-mediated endocytosis, heparan sulfate–dependent surface binding, and endosomal acidification are key factors in exosome internalization by DECs. Conversely, Amiloride, Wortmannin, and Chlorpromazine caused little or no reduction in uptake **(Supplementary Figure 8B),** suggesting that macropinocytosis, phagocytosis, and some dynamin-independent clathrin processes are less important in DEC-mediated exosome internalization. Collectively, these findings show that CTC-exosomes enter maternal decidual stromal cells through specialized, receptor- and acidification-dependent endocytic pathways rather than nonspecific fluid-phase uptake. This mechanistic specificity provides a biological framework for how fetal vesicles deliver transporter cargo to maternal cells in a regulated, energy-dependent manner.

### Exofection restores P-gp expression and efflux function in inflamed and transporter-deficient maternal cells

Having established that maternal DECs efficiently internalize CTC-derived exosomes, we then assessed whether these vesicles can functionally restore P-gp expression and efflux capacity when the transporter is compromised. DECs exposed to LPS (100 ng/mL, 48 h) showed a significant reduction in P-gp protein levels (p < 0.05), consistent with inflammation-induced suppression of ABCB1. Notably, co-treatment with CTC-exosomes (1 × 10¹□ particles) fully restored P-gp expression (p < 0.01), as demonstrated by immunofluorescence analysis **(Figure 6A).** These findings provide direct evidence that fetal vesicles can replenish transporter levels in maternal cells during inflammatory challenges.

**Figure 6:**
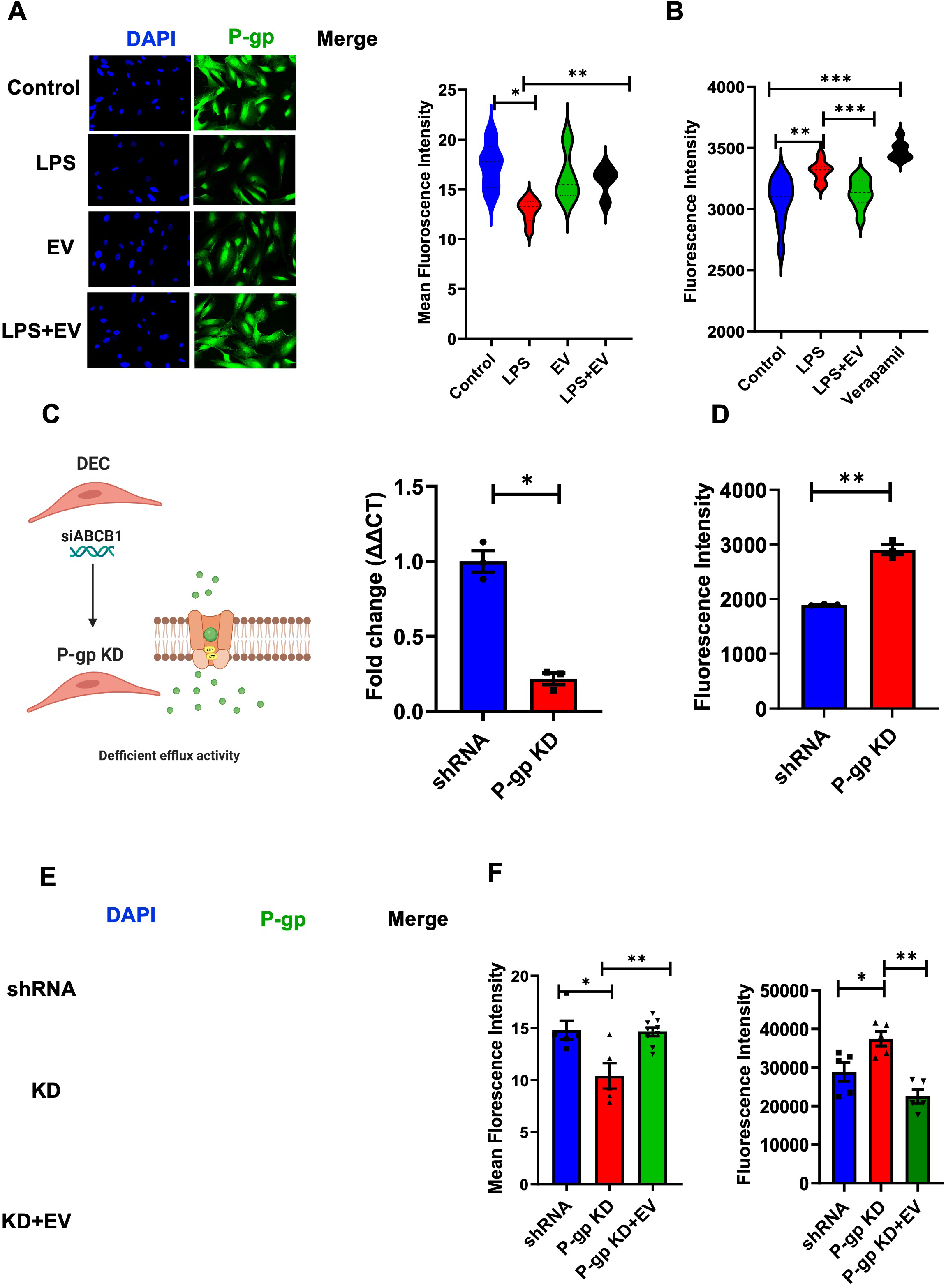
Exofection restores P-gp expression and efflux activity in maternal decidual cells. (A) Immunofluorescence analysis of P-gp expression in DECs under four treatment conditions: Control, LPS, EVs, and LPS + EVs. Nuclei were stained with DAPI (blue) and P-gp with a FITC-conjugated antibody (green). Images captured at 20× magnification. Quantification of P-gp mean fluorescence intensity is shown in the accompanying bar graph. (B) Functional assessment of P-gp activity using a calcein-AM efflux assay across the same conditions. Verapamil was used as a positive control to confirm efflux specificity. (C) Generation of transporter-deficient DECs using ABCB1-targeted siRNA. Knockdown efficiency was validated by qPCR. (D) Loss of P-gp–mediated efflux in knockdown (KD) cells confirmed by increased intracellular calcein accumulation. (E) Rescue of P-gp protein expression in P-gp–deficient DECs following treatment with CTC-derived EVs. Groups include siRNA control, P-gp KD, and P-gp KD + EVs. Representative immunofluorescence images and quantified fluorescence intensities illustrate EV-mediated restoration of P-gp protein. (F) Functional restoration of P-gp activity in KD cells following EV treatment, measured using a dye efflux assay. EV treatment significantly reduced dye accumulation relative to untreated KD cells, demonstrating recovery of transporter function. Statistical significance was determined using Student’s *t*-test or one-way ANOVA. Data are presented as Mean ± SEM. Significance is denoted as * for p < 0.05, ** for p < 0.01 and *** for p < 0.001

To determine if restored protein expression led to improved efflux activity, we performed calcein-AM assays. LPS-stimulated DECs showed a significant increase in intracellular calcein accumulation (p < 0.01), indicating impaired P-gp–mediated efflux. Co-treatment with CTC-exosomes significantly decreased calcein retention (p < 0.01), demonstrating functional recovery of efflux capacity **(Figure 6B).** The addition of the P-gp inhibitor verapamil abolished this effect, causing a marked increase in dye accumulation (p < 0.01), which confirmed that exosome-mediated rescue was transporter-specific.

To determine whether fetal exosomes could restore transporter function in cells lacking endogenous P-gp, we created a transporter-deficient DEC model using ABCB1-targeted siRNA. Knockdown caused a significant decrease in P-gp mRNA (p < 0.05) and a notable increase in intracellular calcein accumulation (p < 0.01), confirming the loss of efflux activity **(Figure 6C–D).** When P-gp–deficient DECs were later treated with CTC-derived exosomes, P-gp protein reappeared in the recipient cells, as shown by restored fluorescence signal and its increased levels **(Figure 6E).** This recovery of protein expression led to functional restoration, as calcein efflux significantly improved after exosome treatment, reducing intracellular dye retention relative to knockdown cells (p < 0.01) and approaching the levels seen in control DECs **(Figure 6F).** Overall, these results demonstrate that fetal CTC-exosomes deliver functional P-gp cargo, capable of restoring transporter expression and efflux capacity in cells deficient either by inflammation or direct gene silencing. These experiments demonstrate exofection as an effective mechanism of intercellular protein transfer and underscore its ability to restore maternal efflux function when transporter loss occurs quickly.

### Exofection restores P-gp–dependent drug clearance and normalizes tacrolimus pharmacokinetics in pregnant P-gp knockout mice

To determine whether fetal exosomes can restore transporter function in vivo, we evaluated the pharmacokinetic consequences of exofection in a pregnant FVB P-gp knockout (KO) mouse model. Animals were assigned to three groups: wild-type (WT), P-gp KO, and KO receiving CTC-derived exosomes (KO+EV). The KO+EV group was administered two intravenous doses of CTC-exosomes (1 × 10¹□ particles; 24 h apart) on gestational day E14 prior to tacrolimus challenge. All animals subsequently received a single dose of tacrolimus (2 mg/kg), a well-characterized P-gp substrate, and plasma levels were quantified by LC–MS/MS across six time points (Figure 7A).

**Figure 7:**
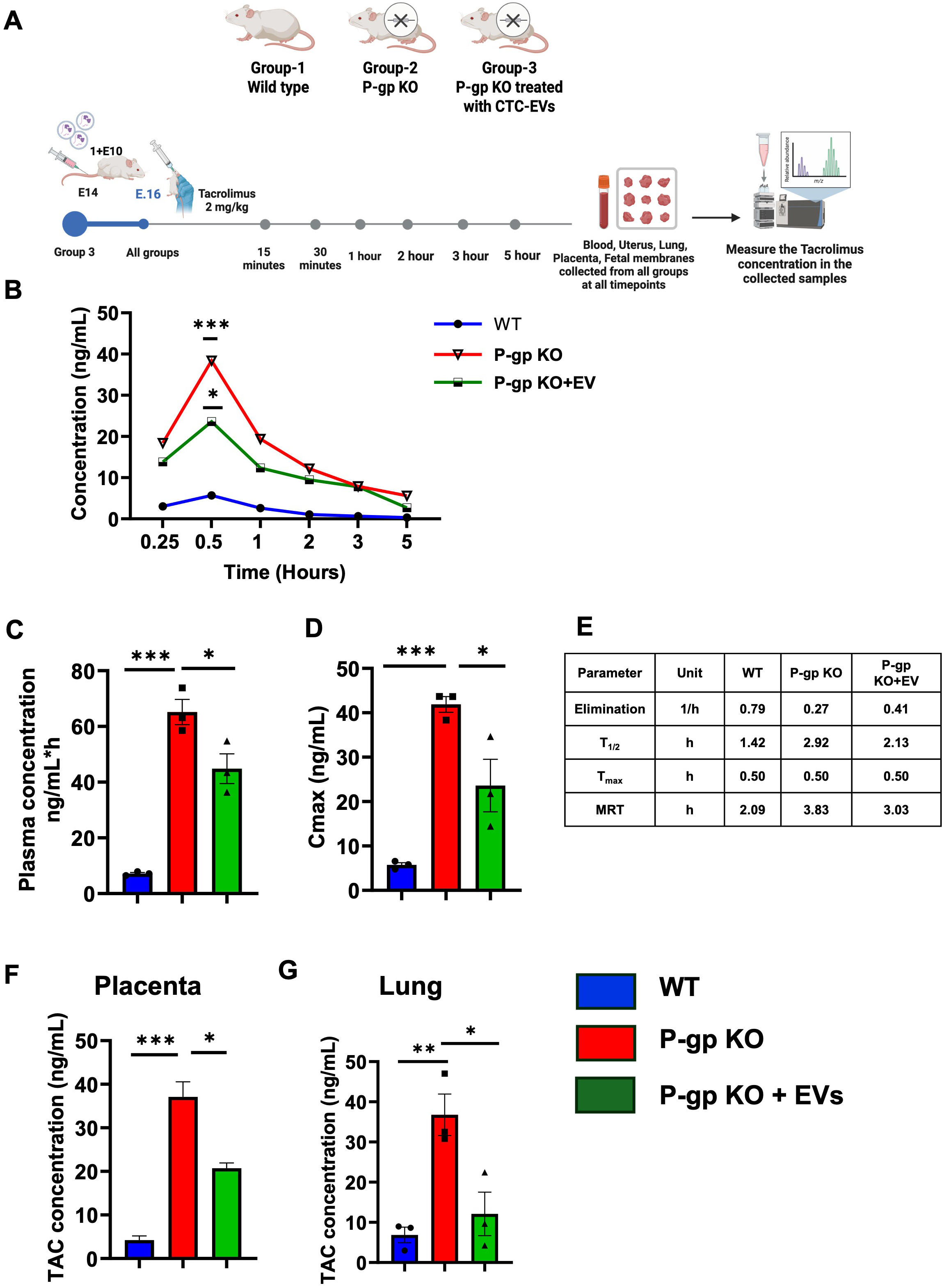
In vivo physiological validation of exofection using pregnant P-gp knockout mice. (A) Schematic overview of the experimental workflow. Pregnant mice were assigned to three groups: wild-type (WT), P-gp knockout (P-gp KO), and P-gp KO treated with CTC-derived exosomes (KO+EV). EVs (1×10¹ particles per dose) were administered on gestational day E14, followed by tacrolimus dosing (2 mg/kg). Plasma and tissues were collected at specified timepoints for pharmacokinetic analysis. (B) Tacrolimus exposure profiles plotted over time for each group. The area under the concentration–time curve (AUC_₀_–_₅_h) was quantified by LC-MS and compared across WT, KO, and KO+EV animals. (C) Tacrolimus plasma concentrations at the 0.5-hour timepoint, illustrating impaired clearance in KO mice and partial restoration following EV treatment. (D) Maximum plasma concentration (Cmax) of tacrolimus in each group. (E) Comparative pharmacokinetic parameters—including elimination rate constant (k_el), half-life (t_₁_/_₂_), Tmax, and mean residence time (MRT)—demonstrating EV-mediated improvement of tacrolimus disposition in P-gp–deficient mice. (F–G) Tissue-specific accumulation of tacrolimus in the placenta (F) and lung (G). EV treatment significantly reduced aberrant drug accumulation in KO animals, indicating partial restoration of P-gp–mediated efflux in vivo. Statistical analyses were performed using Student’s *t*-test or one- or two-way ANOVA where appropriate. Data are presented as Mean ± SEM. Statistical significance is denoted as * for p < 0.05, ** for p < 0.01 and *** for p < 0.001

As expected, P-gp KO mice showed markedly elevated systemic exposure to tacrolimus, with significantly higher plasma concentrations at every time point relative to WT (p < 0.01), reflecting impaired efflux and reduced clearance (Figure 7B). Remarkably, CTC-exosome treatment substantially decreased tacrolimus plasma levels in KO animals, indicating restoration of transporter function following exofection.

Pharmacokinetic analyses reinforced this conclusion. The early exposure metric AUC□–□.□h increased more than eightfold in KO mice (65.19 ng·h/mL) compared to WT (7.9 ng·h/mL), but was significantly reduced in KO+EV mice (44.8 ng·h/mL; p < 0.05) (Figure 7C). Likewise, Cmax rose from 5.7 ng/mL (WT) to 41.8 ng/mL (KO); exosome treatment lowered this value to 23.1 ng/mL (p < 0.05), consistent with restored efflux (Figure 7D).

Assessment of additional pharmacokinetic parameters revealed a similar pattern of rescue. The elimination rate constant (k_el) was substantially diminished in KO animals (0.27 h□¹) compared to WT (0.79 h□¹). CTC-exosome treatment increased k_el to 0.41 h□¹, indicating improved drug clearance (Figure 7E). The mean residence time (MRT), which was markedly prolonged in KO mice (3.83 h), was reduced toward WT levels in KO+EV animals (3.03 h), again supporting partial normalization of tacrolimus disposition. Tacrolimus half-life (t□/□) remained comparable between groups (≈30 min), consistent with P-gp primarily affecting distribution rather than intrinsic metabolism.

Tissue distribution studies revealed profound P-gp-dependent differences in tacrolimus accumulation. KO mice showed significantly elevated drug levels in both lung and placenta (p < 0.001), organs with high physiological reliance on P-gp–mediated efflux. Importantly, CTC-exosome administration reduced tacrolimus burden in both tissues (p < 0.05), indicating that restored transporter function was not restricted to systemic clearance but also re-established protective barrier activity at maternal and fetal interfaces (Figure 7F–G). Reduced tissue retention was further supported by whole-body biodistribution imaging (Supplementary Figure 9). Because selective deletion of P-gp in maternal decidual cells is not yet technically feasible, we used global P-gp KO mice to evaluate whether fetal exosomes can restore systemic P-gp activity in vivo. Although intravenously administered exosomes encounter multiple tissues en route to the uterus, this model directly tests whether fetal exosomes contain functionally competent P-gp capable of rescuing pharmacokinetic defects. The significant improvement in tacrolimus clearance in KO+EV mice supports effective physiological exofection of transporter cargo in vivo.

## DISCUSSION

Exofection(*1*), a novel mechanism identified, was validated in this study using multiple models. Exofection is a process where the donor cell donates cargo through an exosome, the acceptor cell expresses the cargo, where a lost or diminished functional activity is temporarily restored. This is a transient phenomenon in a cell when a specific functional activity from a specific molecule is needed. The FMi cells exchange materials through exofection, and it not only restores the diminished efflux capacity of maternal decidual cells under inflammatory conditions but also re-establishes a protective barrier against xenobiotic accumulation. Using integrated transcriptomic analyses, proteomic characterization, in vitro functional assays, and in vivo pharmacokinetic modeling in P-gp knockout mice, we demonstrate that exofection operates at the FMi as a physiological mechanism of maternal–fetal cooperation. To our knowledge, this is the first demonstration of in vivo functional restoration of a transporter protein by fetal exosomes, revealing a previously unrecognized axis of fetal regulation over maternal physiology.

The restoration of maternal efflux function by the fetus is an indication that fetal membrane cells serve as a robust protective barrier at the FMi even if the maternal system is compromised. This could explain why ascending infections or systemically delivered pollutants, toxicants, pattern-associated markers, and other antigens induced decidual inflammatory changes do not always result in adverse pregnancy outcomes. The higher expression of P-gp in the CTCs was associated with greater efflux activity, suggesting a directional transport mechanism favoring the movement of P-gp’s substrates to the maternal side. These results align with established placental data, where P-gp at syncytiotrophoblast cells similarly facilitates the efflux of compounds into the maternal circulation(*36–39*). In P-gp knockout pregnant mice, we observed impaired clearance of tacrolimus and elevated drug accumulation in maternal tissues(*40, 41*). As hypothesized, treatment with fetal CTC-exosomes restored efflux capacity, evidenced by improved pharmacokinetic parameters, including reduced AUC, lower Cmax, and increased elimination rate in key tissues like the placenta and lungs. Our data provided functional and physiologic validation of *in vitro* experimental evidence for exofection at the FMi in the pregnant animal model.

Exofection occurs in other biological systems and is not restricted to pregnancy. Healthy brain endothelial cells (BECs) donate mitochondria and HSP27 via medium-to-large exosomes to ischemic BECs, restoring ATP production, mitochondrial function, and tight junction integrity(*4*). Similarly, TNF-α-preconditioned human umbilical cord mesenchymal stromal cells deliver exosomes containing 3,4-dihydroxyphenylglycol to acinar cells affected by severe acute pancreatitis, reestablishing cellular homeostasis through mTOR pathway activation, which inhibits autophagy, inflammation, and oxidative stress(*2*). Similarly, in early pregnancy, first-trimester placental cells secrete exosomes that induce programmed death-ligand 1 (PD-L1) expression in maternal monocytes via the PTEN pathway, thereby enhancing maternal immune tolerance toward the fetus(*42*). Our data support the notion that the fetus is not merely a passive recipient of maternal regulation but an active participant in modulating maternal physiology through targeted vesicular communication. Exofection at the FMi occurs via heparan sulfate proteoglycan-mediated (HSPG), dynamin-dependent endocytosis and pH-dependent endosomal fusion, which are well-established exosome uptake mechanisms(*43–48*). Targeting any of these pathways helps to enhance exosome uptake. One such work is neural stem cell-derived exosomes, which utilize HSPG-mediated endocytosis (HSPG as a receptor for EVs) to cross the blood-brain barrier, offering a promising nanocarrier platform for brain-targeted drug delivery(*49*).

Furthermore, exosome-mediated transfer of functional proteins is well established in pregnancy. Our lab has established this exosome-mediated paracrine signaling at FMi(*50–55*). While past studies have proposed fetal exosomes as immunomodulators during pregnancy(*56, 57*), this work identifies their capacity to restore barrier function at the FMi through transporter protein-specific exofection. This reveals a fetal-initiated, compensatory system that safeguards fetal health when maternal clearance pathways are impaired. Notably, enrichment of different TP families suggests active involvement in solute and molecule exchange. These features underscore the potential of CTC exosomes to mediate important communication at the FMi and highlight their role as biological interventions for the delivery of therapeutics in pregnancy-related disorders.

Despite providing strong evidence that exofection serves as a mechanism of material transport and cellular engineering at the feto-maternal interface, this study has certain limitations. While we demonstrate that exosomal delivery of P-gp can restore efflux function in maternal decidual cells, we have not yet established whether the absence of exofection directly contributes to adverse pregnancy outcomes, such as preterm birth. Moreover, although P-gp is a critical efflux transporter, its precise role in the pathogenesis of pregnancy complications remains to be elucidated. Ongoing studies in our lab are addressing this gap using ascending infection-induced preterm birth models in P-gp knockout mice, which will allow us to assess the physiological consequences of impaired transporter function and the potential protective role of exofection.

Additionally, this study focused primarily on the transfer of protein cargo; however, from our supplementary data suggest that RNA molecules may also be exofected. Notably, early time-point analyses revealed changes in gene expression in the DECs following CTC-exosome treatment, which upregulates ABCG1 in DECs, suggesting possible exosomal RNA-mediated gene modulation. While these findings are intriguing, further work is needed to determine whether RNA exofection leads to stable expression and functional protein translation, and whether these effects are transient or sustained. We are currently developing a CRISPR-based maternal-specific P-gp knockdown model, sparing fetal expression, to dissect the directional role of exofection in vivo. Collectively, these limitations underscore the need for more refined genetic tools and longitudinal functional studies to elucidate the biological and therapeutic implications of exofection fully.

Together, exofection has a potential role in pharmacology in reversing the pathologic phenotype of a cell that has lost or diminished the capacity of an indispensable protein. Exofection of a desired cargo, using engineered exosomes that carry these substances, can therapeutically benefit the cell. The exosome cargo-mediated cellular engineering process enables the cell to acquire a functional capability to reverse its pathologic phenotype to a normal one. Based on our observation, exofection is a transient phenomenon that does not permanently alter the cells’ functions; therefore, the long-term consequences of exofection processes can also be minimal. Unlike other secreted factors (e.g., cytokines) that can continually induce functional changes, exosomes are likely to perform these functions *in vivo* to ensure the transient nature of exofection and to minimize long-term implications. The properties of exosomes, their biogenesis by the donor cell, cargo packaging efficiency, release, and targeting of recipient cells are not trivial. Exofection of the recipient cells tends to be based on the need and depends on the environment (e.g., inflammation at the FMi). The recipient cell should also possess the necessary cellular machinery to express the specific cargo and perform its intended function. Endo-lysosomal processes also eliminate exosomes, and a constitutive expression and function of the incoming cargo is unlikely. Therapeutically, this can be beneficial in transitioning to a cellular phenotype or achieving a functional capability. Multiple approaches to engineering exosomes have now been developed and tested for therapeutic purposes. Exofection of a lost protein, mRNA or other molecule should be a therapeutic option for many diseases.

## CONCLUSION

In conclusion, our study uncovers exofection as a novel, physiologically relevant mechanism of intercellular transport at the FMi, wherein fetal-derived exosomes deliver functional transporter proteins to maternal cells. Through integrated transcriptomic, proteomic, *in vitro*, and *in vivo* analyses, we demonstrate that this process restores efflux capacity and maintains barrier integrity under inflammatory conditions and transporter deficiency. These findings redefine fetal-maternal communication as a bidirectional, compensatory system, advancing our understanding of pregnancy homeostasis. Beyond pregnancy, exofection represents a broader biological principle with implications for regenerative medicine, drug delivery, and the treatment of diseases characterized by acute or reversible protein loss. As engineered exosome technologies continue to advance, exofection may emerge as a versatile therapeutic approach capable of temporarily reprogramming dysfunctional cells without permanent genetic modification.

## MATERIALS AND METHODS

### Sample collection and processing

The UTMB Institutional Review Board (IRB) in Galveston, TX, approved the collection of human placentas under protocol UTMB 11-251. These discarded placentas, obtained from cesarean deliveries without labor, were collected at John Sealy Hospital. Following the approved inclusion and exclusion criteria, our team harvested and processed the placentas and fetal membranes using a previously established methodology(*58, 59*). In summary, the fetal membranes were separated from the placentas, and samples were excised from the central region of the membranes. These samples were then processed as explants for further analysis. The study utilized anonymous discarded placental specimens, which did not qualify as human subjects under the Health and Human Services regulations outlined in 45 CFR part 46.102.

### Cell culture

Fetal membrane CTCs were obtained from term individuals, not in labor (TNIL), and immortalized using the HPV16 E6E7 retrovirus method(*58, 60*). CTCs were cultured in a specialized DMEM/Ham’s F12 medium containing penicillin, streptomycin, bovine serum albumin, heat-inactivated fetal bovine serum (HI-FBS), and ITS-X, along with CHIR99021, A83-01, SB431542, L-ascorbic acid, valproic acid (VPA), Y27632, and epidermal growth factor to support their proliferation and maintenance. Primary DECs were cultured in Dulbecco’s Modified Eagle Medium (DMEM) and Ham’s F12 nutrient medium (1:1), supplemented with antibiotics (amphotericin B, penicillin, and streptomycin), 10% HI-FBS. These cells were used up to the 20th passage to ensure experimental consistency. All cultures were maintained in a standard incubator at 37°C with 5% CO□ and 95% humidity, using the method of isolation mentioned in the previous publications(*58, 60*).

### Immunohistochemistry

Fetal membrane tissue sections were obtained from the previously isolated TNIL placenta. After 48 hours of formalin fixation, tissues were transferred to 70% ethanol, followed by dehydration and paraffin embedding through graded ethanol, xylene, and paraffin steps. Paraffin-embedded tissues were sectioned at a thickness of 5 μm. Sections were deparaffinized overnight at 50°C, washed with xylene and ethanol, and subjected to antigen retrieval by boiling in Tris-EDTA buffer (pH 9.0; Abcam, Cat# Ab93684) for 2 hours. The peroxidase activity was quenched by incubating sections with 3% hydrogen peroxide for 10 minutes. This was followed by a 5-minute blocking step using a blocking solution from Sigma Millipore (Cat# 4061341), and slides were washed three times for 5 minutes each. Sections were then incubated for 20 minutes in a humidified chamber with an anti-human primary antibody against P-gp (Abcam, Cat# ab235954) negative control sections, in which no primary antibody control, were processed in parallel and. After washing, incubated with a secondary antibody from the kit for 10 minutes, followed by additional washes. Two drops of streptavidin reagent were then applied, and sections were incubated for 10 minutes in a humidified chamber. Following a wash step, chromogen reagent (A+B) was added for 10 minutes and rinsed as previously described. Counterstaining was performed using hematoxylin for 1 minute, followed by a washing step. Slides were dehydrated using different alcohols and xylenes and then mounted using a xylene-based mounting medium (Acrymount). Slides were examined by bright-field microscopy, and images were captured at 10X magnification.

### RNA isolation and Quantitative PCR

CTCs and DECs cell pellets were homogenized with 350 μL RNA lysis buffer, followed by adding one volume of 70% ethanol to the homogenate. This mixture was transferred to RNeasy spin columns at a speed of 12,000 *g* for 30 seconds at 4°C. After discarding the flow-through, RNA wash buffer was added to the columns, which were then centrifuged again under the same conditions. The columns were washed twice with RW1 wash buffer, and RNA was eluted with RNase-free water by spinning at 12,000 *g* for 1 minute. RNA yield and purity were assessed using ultraviolet-visible spectrophotometry, with all samples showing A260/A280 values between 1.8 and 2.0. Total RNA (1 μg) was reverse-transcribed using a high-capacity RNA-to-cDNA kit (Applied Biosystems). qPCR reactions were performed using taqman™ PCR master mix (Applied Biosystems) and taqman™ gene expression assay (FAM) probes for ABCB1 HS00184500 (Life Technologies) with GAPDH (Life Technologies, Hs00177504) as internal control. Gene expressions were calculated.

### Flow cytometry

CTCs and DECs were detached using Accutase (Thermo-Fisher Cat# 13150016) at 37°C for 8 minutes. Following detachment, cells were washed with PBS and resuspended in fixation and permeabilization solution (Thermo-Fisher Cat# 00-5523-00). The cells were incubated for 20 minutes at 4°C. Followed by washing with 2□mL of permeabilization buffer and then blocked using Fc block (Cat# 564220, 1:100 dilution in permeabilization buffer) for 30 minutes in the dark at room temperature. Cells were rewashed and centrifuged at 1500 rpm for 5 minutes. The cells were resuspended in 100□µL of FACS buffer (PBS+2% FBS), and 4□µL (ABCB1-PE, Lot: B330626) of the P-gp intracellular antibody was added and incubated for 45 minutes at room temperature in the dark. Following incubation, cells were washed at 1500 rpm for 8 minutes and resuspended in 200□µL of FACS buffer for analysis. An unstained sample was included as a control. The samples were analyzed using a CytoFLEX flow cytometer, and data were analyzed using FlowJo software.

### Transwell experiment for testing fetal membrane/decidua efflux activity

Fetal membrane explants (2 × 2 cm) were prepared from freshly collected TNIL placentas. The placental tissues were thoroughly washed with sterile saline three times to remove blood and debris. Corning transwell inserts without the membrane (Cat# 07-200-161) were used for mounting the explants. The fetal and maternal sides of the tissue were separated, and mounting was initiated using sterile rubber bands. To ensure proper attachment, a small, rough surface was created using scalpels on the trans well walls. After securing the tissue to the insert, the integrity of the mounted membrane was assessed by adding culture media to the upper chamber and checking for any leaks into the lower chamber. Only wells without leakage were used for further analysis. Selected Transwells were filled with media on the apical (top) and basal (bottom) sides and incubated for 24 hours at 37°C. Tissue viability was assessed based on the change in the media color to yellow. Upon confirmation of tissue integrity, the media were replaced with fresh media containing 80 µg of tacrolimus on the apical side. After a 3-hour incubation, media samples were collected from the basal side of the wells and stored for LC-MS analysis to quantify tacrolimus.

### Transwell experiment for efflux activity-cells

FMi cells, DECs and CTCs were seeded onto Transwell membranes (Corning, Cat# 07-200-161) at a density of 500,000 cells/mL and incubated at 37°C with 5% CO□ until a confluent monolayer was observed. The inserts featured a polyester membrane with a 0.4 μm pore size. Media were replaced as needed to maintain the cell culture. Upon confirmation of monolayer formation, the culture media were replaced with fresh media containing 40 µg of tacrolimus, added to the apical side of the transwell. Media samples were collected from the basal side after a 3-hour incubation period and stored for subsequent LC-MS analysis.

### Cells and Exosome Transcriptomics

#### Sequencing library preparation

RNA samples were quality-checked using an Agilent bioanalyzer RNA nanochip. RNA sequence libraries were prepared using NEB Next PolyA module (Cat# E7490, NEB, Ipswich, MA) combined with NEB Next Ultra II RNA Library preparation kit (Cat# E7775, NEB, Ipswich, MA) following the manufacturer’s recommended procedure. The resulting libraries were quality-checked using an Agilent bioanalyzer HS DNA chip. The libraries were pooled and sequenced on ElementBio (San Diego, CA) Aviti sequencer for PE150 sequence targeting approximately 80 million reads per sample. Sequencing data analysis: Sequencing reads were quality-checked using fastqc (v.0.12.1, Andrews, S. 2010), and then adapter sequences were trimmed from the reads using trimmomatics (v.0.39, Bolger et al., 2014). The trimmed reads were mapped to human genome hg38 using STAR (v2.7.11a. Dobin et al., 2013) and differential expression analyzed using Bioconductor (v.3.19.1. Gentleman et al. 2004) and DESeq2 package (v. 1.44.0. Love et al, 2014). GO enrichment test was performed using clusterProfiler. For the EV samples, the ribosomal RNA was depleted using the NEBNext rRNA depletion kit (E7400) instead of polyA enrichment, and the rest is the same as the cell RNA library preparation.

### CTC-EV proteomics

Analyzed peptide mixtures using nanoflow liquid chromatography-tandem mass spectrometry (nanoLC-MS/MS) on an UltiMate 3000 RSLCnano system (Dionex) coupled online to a Thermo Orbitrap Eclipse mass spectrometer via a nanospray ion source. Peptides were directly injected onto a C18 analytical column (Aurora, 75 µm × 25 cm, 1.6 µm; IonOpticks) after equilibrating in 98% solvent A (0.1% formic acid in water) and 2% solvent B (0.1% formic acid in acetonitrile). We injected 2 µL of sample at 300 nL/min and applied a multistep gradient for elution. Data acquisition was performed in positive ion mode using data-independent acquisition (DIA) with staggered 8 Da windows (400–900 m/z), a 3-second cycle time, and Orbitrap detection at 60,000 resolution for MS1 and 30,000 for MS2. We demultiplexed raw data to mzML using MSConvert and analyzed them in Fragpipe v22.0 with MSFragger for database searching and DIA-NN for quantification. Search parameters included trypsin/P digestion (up to 2 missed cleavages), fixed cysteine carbamidomethylation, variable methionine oxidation, N-terminal acetylation, and 1% precursor FDR. The Homo sapiens UniProt database used (UP000005640, updated 2025-02-05) for protein identification. Filtered out the contaminants, single-peptide hits, and inconsistently quantified proteins, retaining those with ≥20% non-missing values globally and ≥50% in at least one condition. Data were log2-transformed, variance stabilized and analyzed using **limma** in R for differential expression. Proteins with an adjusted p-value < 0.05 and absolute log2 fold change ≥1 were considered significantly regulated.

### EV isolation by the tangential flow filtration (TFF) system

Our team optimized the isolation and purification of EVs using the TFF protocol. Culture media from CTCs at 70-80% confluency were collected and stored at -80°C. To pre-clean the media before EV isolation, sequential centrifugation was performed: 300 *g* for 5 minutes to remove cell contamination, 3000 *g* for 10 minutes to eliminate cell debris, and 17,000 *g* for 15 minutes to discard microvesicles, followed by filtration through a 0.2 µm membrane to retain only particles smaller than 0.2 µm. The TFF system was then prepared with an exosome membrane module. The system was first flushed with water through both the retentate and filtrate tubes, sterilized with 0.5M NaOH, and rinsed with water. The pre-cleaned media was introduced into the sample chamber, where flow rate (30-40 mL/min), pressure (20-30 psi), and room temperature were carefully maintained to ensure stable filtration. The TFF 100 K membrane filtrate effectively removed particles smaller than 100 kDa, including free soluble proteins. Finally, the purified EV sample was collected and further concentrated into a small volume using Beckman ultracentrifugation.

### Exosome size distribution and concentration measurement by Zetaview

Exosomes concentration and size distribution were analyzed with the ZetaView™ PMX 110 (Particle Metrix, Meerbusch, Germany) with 8.02.28 software. The Zetaview machine was prepared by washing the main line and sample injection line with water (3X) before use. The instrument calibration was done with the beads at a 1:250000 dilution. Frozen Exosome samples were gently thawed on ice and then diluted in filtered water at a 1:1000 ratio. This dilution allowed for accurate measurement of particle concentration (particles/mL) and average size distribution for each sample. To ensure the instrument’s accuracy, it was thoroughly rinsed with filtered distilled water in between each sample analysis.

### Exosome tetraspanin marker identification by Nanoview imager

The isolated Exosomes were immobilized on poly-L-lysine-coated coverslips. Fix Exosomes with 4% paraformaldehyde (PFA) were immunolabeled with antibodies against CD9, CD63, and CD81, followed by Alexa Fluor 647-conjugated secondary antibodies. Super-resolution imaging was performed using the ONI Nanoimager S with a 100x oil immersion objective and 640 nm excitation. Localization-based imaging (dSTORM) was conducted over 15,000 frames using ONI imaging buffer. Image reconstruction, drift correction, and particle analysis were carried out using ONI’s CODI software to assess exosome size, count, and marker co-localization.

### Exosome P-gp quantification by ELISA

Prior to beginning the experiment, all reagents (Human P-gp/ABCB1 ELISA kit, Cat# RDR-Pgp-Hu from Reddot) were allowed to equilibrate to room temperature. Then, 100 μL of the standards and Exosome samples (protein concentration measured by BCA) were added to the designated wells, which were sealed and incubated at 4°C overnight. In the morning, the wells were washed three times for 2 minutes with 350 μL of wash buffer. Next, 1X 100 μL of biotinylated detection antibody A was added to the wells, and they were incubated at 37°C for 60 minutes. The wells were then washed five times with wash buffer. Following this, 1X 100 μL of HRP-streptavidin conjugated detection antibody B was added, and the wells were incubated at 37°C for 60 minutes. After another series of five washes with wash buffer, 90 μL of substrate was added, and the wells were incubated at 37°C in the dark for 10-20 minutes. 50 μL of stop solution was added to each well, and the optical density (O.D. value) was immediately measured at 450 nm using a microplate reader.

### P-gp and tetraspanin marker characterization by western blot analysis

CTC-EVs were lysed using 10X radioimmunoprecipitation assay buffer containing 50 mM Tris (pH 8.0), 1% Triton X-100, 1.0 mM EDTA (pH 8.0), 150 mM NaCl, and 0.1% SDS, supplemented with 1X phenylmethylsulfonyl fluoride, and 1X protease and phosphatase inhibitors. Following centrifugation at 12,000 *g* for 20 minutes, the supernatant was collected, and the protein amount was calculated using the Pierce BCA protein assay kit (Thermo Scientific, Waltham, MA, USA). Protein samples were run via SDS-polyacrylamide gel electrophoresis (10%) and transferred to a membrane using a Bio-Rad gel transfer system. The membranes were cut and blocked for 2 hours at room temperature (shaking) with 5% milk in 1X Tris-Buffered Saline, 0.1% Tween (TBST), and then incubated overnight at 4°C with primary antibodies (shaking). The membranes were washed four times and incubated for 1 hour with secondary antibodies, and protein bands were visualized using a chemiluminescence (ECL max) detection system (Amersham, Piscataway, NJ, USA). Primary antibodies included EV markers CD63, CD81, and CD9 (1:1000, SBI Systems Biosciences, Cat# EXOAB-kit-1, Lot# 30227-003), P-gp (1:1000, Abcam, Cat# ab170904). Secondary antibody for P-gp anti-rabbit (1:10,000, Amersham, Cat# NA934VS), and for the CD markers, the secondary antibodies were provided by the SBI kit EXOAB-kit-1 at a 1:5000 dilution.

### Automated Western Blot Analysis

Exosome samples were prepared by diluting 1:20 in RIPA lysis buffer (50 mM Tris pH 8.0, 150 mM NaCl, 1% Triton X-100, 1.0 mM EDTA pH 8.0, 0.1% sodium dodecyl sulfate), supplemented with protease and phosphatase inhibitor cocktail and phenylmethylsulphonyl fluoride. Protein concentration was quantified using the Pierce BCA assay (Thermo Fisher Scientific). Automated western blotting was then performed using a JESS system (Protein Simple, California, USA) following the manufacturer’s protocol. For analysis, 3 µL of each sample was diluted with 0.1X Sample Buffer (containing SDS, DTT, and fluorescent molecular weight standards) to a final protein concentration of 0.3 mg/mL. Samples were denatured at 70 °C for 10 min and then loaded into the plate. Default settings were applied for the run in 25 capillary cartridges (12-230 kDa, Cat no. SM-W004, Bio-Techne, Minneapolis, USA). Primary antibodies for exosome markers—FLOT-1 (Cat no. 18634S, Cell Signaling), CD9 (Cat no. 13174S, Cell Signaling), CD81 (Cat no. Exoab-CD81A1, System Biosciences), CD63 (Cat no. 52090S, Cell Signaling), and Calnexin (Cat no. 2433S, Cell Signaling). Peaks and bands were subsequently detected and analyzed using Compass for Simple Western software (Version 6.3, Bio-Techne).

### Cryogenic electron microscopy (cryo-EM)

The exosome samples were thawed and added to glow-discharged Quantifoil holey carbon grids (Cu 300 mesh, R1.2/1.3), blotted for 2–3 seconds under controlled humidity (∼100%) and temperature (4°C), and rapidly plunge-frozen in liquid ethane using a Vitrobot Mark IV (Thermo Fisher Scientific). Grids were transferred to a Titan Krios transmission electron microscope (Thermo Fisher Scientific) operating at 300□kV and imaged using a direct electron detector (e.g., Gatan K3 or Falcon 4). Movies were recorded in counting mode at a nominal magnification of ×81,000, yielding a calibrated pixel size of ∼1.1□Å. Data were collected using automated acquisition software (e.g., EPU or SerialEM) with a total electron dose of ∼50□e□/Å² distributed over 40 frames. Motion correction, contrast transfer function (CTF) estimation, particle picking, 2D classification, and 3D reconstruction were performed using RELION or cryoSPARC software suites.

### Lactate dehydrogenase (LDH) assays for EV uptake inhibitors

Cell supernatants were collected from drug-treated wells after 24 hours. According to the manufacturer’s protocol, 5□μL of each supernatant was mixed with 95□μL of LDH reagent (ab197004, Abcam, Cambridge, UK) in a 96-well plate and incubated in the dark at room temperature for 10 minutes. Fluorescence was measured immediately using a microplate reader (excitation: 535□nm, emission: 587□nm). Fresh culture medium (5□μL) served as a blank. To establish a positive control for 100% cytotoxicity, cell supernatants were treated with 10 μL of the kit’s cell lysis buffer to induce complete membrane lysis.

### EV PKH-26 staining and inhibitor treatment for the maternal EV uptake studies

To evaluate the propagation of exosomes across maternal DECs, exosomes were labeled using PKH-26 dye (PKH26PCL, Sigma, MKCS0369). Briefly, 4□μL of PKH-26 was diluted in 1□mL of Diluent C. After a 5-minute incubation; the reaction was quenched with 1% BSA. Unbound dye was removed by 30 minutes of centrifugation at 700 *g* using Amicon 100 K filters: samples were centrifuged two times and transferred to fresh filters. The labeled exosomes were then washed twice with PBS, centrifuged at 1000□×□*g* for 10 minutes between washes, and the flow-through was discarded. The labeled exosomes were introduced into the maternal decidua, and their propagation was assessed. For the inhibitor studies, drugs such as Dynasore (Abcam, Cat# ab120192), Wortmannin (Abcam, Cat# ab120148), Amiloride (Abcam, Cat# ab120281), Chlorpromazine (Sigma, Cat# 31679), Omeprazole (Sigma Cat# PHR1059), and Heparin (Sigma Cat# H3393) were added to the cells before 1 hour and treated with PKH-26-labeled exosomes for 4 hours.

### Treatment of DECs with LPS and CTC exosomes

DECs and CTCs were seeded at a density of 0.20 × 10□ cells per well in six-well plates and cultured for 70–80% confluency. For the inflammatory model, DECs and CTCs were treated with 100 ng/mL LPS (LPS; Sigma, Cat# L2880), prepared from a 1 mg/mL stock solution. Cells were exposed to LPS for 48 hours to induce inflammation. For exofection experiments, cells were co-treated with 100 ng/mL LPS treatment and exosomes at a concentration of 1 × 10¹□ particles for 48 hours. The LPS concentration of 100 ng/mL was selected based on prior literature(*61*) and experimental optimization, ensuring sufficient induction of an inflammatory response without compromising cell viability.

### Analysis of inflammatory cytokines by multiplex assay

Culture media from both cell types were collected 48 hours post-treatment to assess the inflammatory response. Levels of IL-6, IL-8, and GM-CSF were quantified using a Luminex-based multiplex assay (HCYTA-60K, EMD Millipore, Burlington, USA). The assay kits demonstrate high precision, with intra-assay variability of less than 15% and inter-assay variability below 20%, and an accuracy range of 92%–106%. Standard curves were established using duplicate samples of recombinant cytokines with known concentrations, as provided by the manufacturer. Briefly, standards and samples were incubated with magnetic capture beads conjugated to specific antibodies in a 96-well plate overnight at 4°C on a shaker. Following washes, biotinylated detection antibodies were added and incubated for 30 minutes, followed by the addition of streptavidin-phycoerythrin (PE) for fluorescent signal development. Cytokine concentrations were analyzed from standard curves using a 5-parameter logistic regression model in Bio-Plex Manager™ software.

### Immunocytochemistry

After the LPS treatment, the cells were first fixed and permeabilized with 4% PFA and 0.5% Triton X and blocked with 3% BSA in PBS. The cells were then incubated overnight at 4°C with a primary antibody targeting P-gp (Abcam, Cat# ab235954), diluted at 1:300. Following thorough PBS washes, the cells were incubated for 1 hour with a secondary antibody, Alexa Fluor 488-conjugated anti-rabbit IgG (Abcam, Cat# ab150073), at a dilution of 1:1000. After additional PBS washes, the cells were stained with NucBlue™ Fixed Cell Ready Probes™ reagent (Invitrogen) and then mounted using Mowiol (Calbiochem Cat# 475904) mounting medium.

### P-gp efflux assay

The efflux dye assay was performed using the Multidrug Assay Kit (Cayman Chemicals, item # 600370) according to the manufacturer’s protocol. The treated cells were washed with PBS and incubated at 37°C for 30 minutes with 100 μL media added with or without Verapamil (P-gp inhibitor at 1:2000 dilution), followed by an additional 30-minute incubation at 37°C with 100 μL calcein dye (2 μL in 10 mL media). After the incubation, we washed the media by centrifugation at 400 *g* for 5 minutes, twice with cold media, removed the media containing dye, added 200 μL of fresh cold media, and measured the fluorescence intensity at 485 excitation and 535 nm emission wavelength.

### DECs P-gp knockdown

Knockdown experiments were performed in DECs using P-gp-targeted small interfering RNAs (siRNAs). Briefly, siRNA at a concentration of 30 pmol for each target was delivered to DECs at 80% confluence per well of a 6-well plate via transfection using DharmaFECT1 transfection reagent. For the knockdown control, the cells were transfected with non-targeting control siRNAs. The transfected cells were incubated for 48 hours prior to confirmation of knockdown and subsequent efflux assay. The knockdown confirmation was achieved by RT-qPCR analysis using reverse-transcribed RNA samples isolated from siRNA-transfected DEC. Upon confirmation of knockdown, the cells were transferred into a 96-well plate from which efflux activity was analyzed after 24 hours. The efflux activity was determined using a fluorometric multidrug resistance (MDR) assay (AAT Bioquest), where the impact of knockdown on efflux activity was determined by fluorescent intensity detected in the cells.

### Animal studies

Animal procedures were followed in accordance with the Institutional Animal Care and Use Committee (IACUC) at the University of Texas Medical Branch, Galveston, under approved protocol number 041107F. Timed-pregnant FVB/NTac wildtype mice (Taconic Biosciences) were obtained on gestational day E.13. The mice were housed in a temperature-controlled facility. For the P-gp KO, FVB.129P2-Abcb1a Abcb1b N12, females and males were purchased and bred in the UTMB animal facility. Pregnancy was confirmed by identification of a sperm plug and subsequent weight gain. The animals were divided into three groups: WT, KO, and KO+EV. On E.14, KO+EV group mice were treated with two doses of CTC-exosomes (1 × 10¹□) every 24 hours for 48 hours. On E.16, all three groups received tacrolimus 2 mg/kg, chosen based on literature and pharmacokinetics of tacrolimus drug for 15 minutes, 30 minutes, 1 hour, 2 hours, 3 hours, and 5 hours(*41*). After completion of the time points, the mice were sacrificed, and the blood, lungs, placenta, fetal membrane, and uterus were isolated for further analysis.

### Mouse tissue processing

After the experiment, the isolated tissue was frozen. Then, the tissue was thawed, and the samples were weighed on the glass slide. They were then kept at -80°C for 30 minutes. Then, the tissue was finely cut, and PBS was added in a 1:4 ratio. Finally, half a spoonful of 1mm Zrsio beads was added. Using the bullet blender, the tissue was homogenized until no tissue remained. The sample was collected and stored at -80°C.

### Analytical method for Tacrolimus measurement

A liquid chromatography-tandem mass spectrometry (LC-MS/MS) method with electrospray ionization (ESI) was developed and validated for quantifying tacrolimus in mouse plasma, placenta, fetal membrane, uterus, and lung samples following FDA guidelines. Tacrolimus-*d3* was selected as the internal standard. The chromatographic separation of tacrolimus and its internal standard was performed on a Waters XTerra® MS C18 HPLC column (50 × 2.1 mm, 3.5 µm) at 45°C. The mobile phase consisted of acetonitrile + water with 2% NH□OH (v/v), and gradient elution was used at a flow rate of 350 µL/min. An API 4000 triple quadrupole mass spectrometer, equipped with a Turbo V ion source, operated in negative mode. Quantification was conducted using multiple reaction monitoring (MRM) with transitions of m/z 802.5 → 560.6 for tacrolimus and m/z 805.5 → 563.6 for tacrolimus-*d3*. The calibration range for tacrolimus in mouse serum ranged from 0.26 ng/mL to 44.8 ng/mL. Method accuracy ranged from 92% to 112%, with relative standard deviation (RSD) values not exceeding 11%. Quality control samples at high, medium, and low concentrations were analyzed alongside test samples to ensure result reliability.

### Statistical analysis

Data were analyzed as mean values with the standard error of the mean (SEM) indicated. Two-tailed Student’s t-tests were utilized to compare the two groups. When the analysis involved more than two groups, one-way ANOVA was performed, employing a general linear model with a univariate analysis approach. More than three groups were analyzed using two-way ANOVA with multiple comparisons. All the experiments were performed with three biological replicates or three different experiments.

### Softwares

GraphPad Prism 10 (www.graphpad.com) was used for data plotting and statistical analyses. A p-value less than 0.05 was considered statistically significant. BioRender was used to create schemes illustrating the experimental model and the graphical representation.

## List of Supplementary Materials

Figure S1 : Flow cytometry gating strategy for DEC with unstained control for P-gp-PE

Figure S2 : Flow cytometry gating strategy for CTC with unstained control for P-gp-PE

Figure S3 : IPA analysis of the subcellular localization of gene networks associated with ABCB1 downregulation in DECs

Figure S4: IPA analysis of the subcellular localization of gene networks associated with ABCB1 upregulation in CTCs

Figure S5 : Morphological identification of CTC exosomes using Cryo-EM

Figure S6 : Exosomes characterization by simple Western Jess (automated western system)

Figure S7 : Gene ontology enrichment analysis generated by Metascape

Figure S8 : Mechanisms of maternal DEC cell uptake of EVs

Figure S9 : Drug elimination efficiency of CTC exosomes with P-gp as a cargo

## Acknowledgments

We sincerely thank **Ms. Rheanna Urrabaz-Garza** for her invaluable assistance in securing the animal amendment in a timely manner and **Ms. Phyllis Gamble** for her dedication to maintaining the animal colony. These individuals’ contributions were instrumental in the successful completion of this study.

## Author contributions

Conceptualization: AK, RM

Methodology: MT, AW, VM, PE, EA, AK, RM, LSR, MT

Investigation: MT, AW, TJT, XMW

Visualization: LR, AK

Funding acquisition: AK, RM

Project administration: AK, RM

Supervision: AK, LR, RM

Writing – original draft: MT,AK, RM,

Writing – review & editing: VM, PE, EA,LR

## Data and materials availability

All data are available in the main text or the supplementary materials.

