## Supplementary figures and images for "P-glycoprotein exofection between fetal and maternal cells as a mechanism of intercellular material transfer at the feto maternal interface"

### Supplementary figure 1

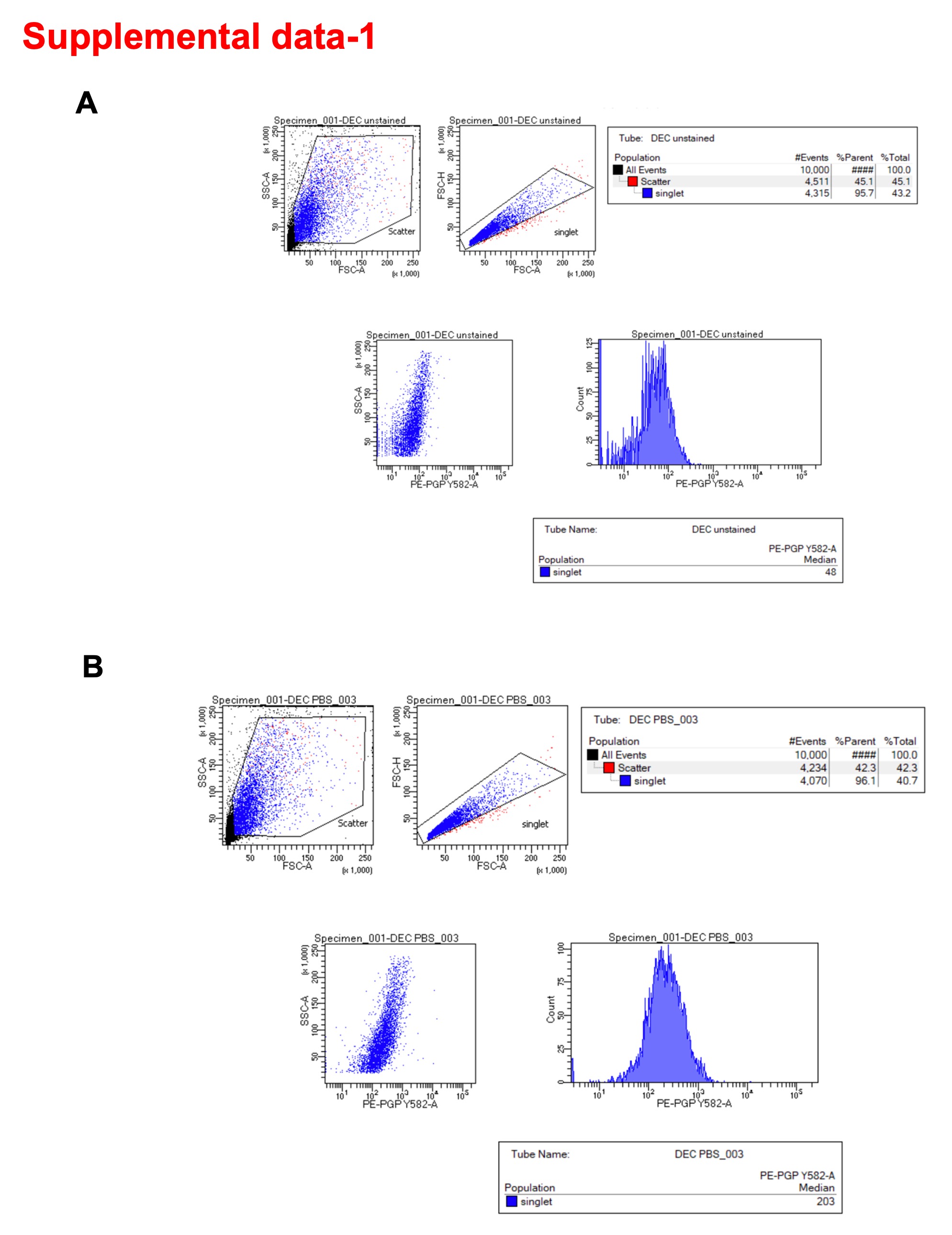

### Supplementary figure 2

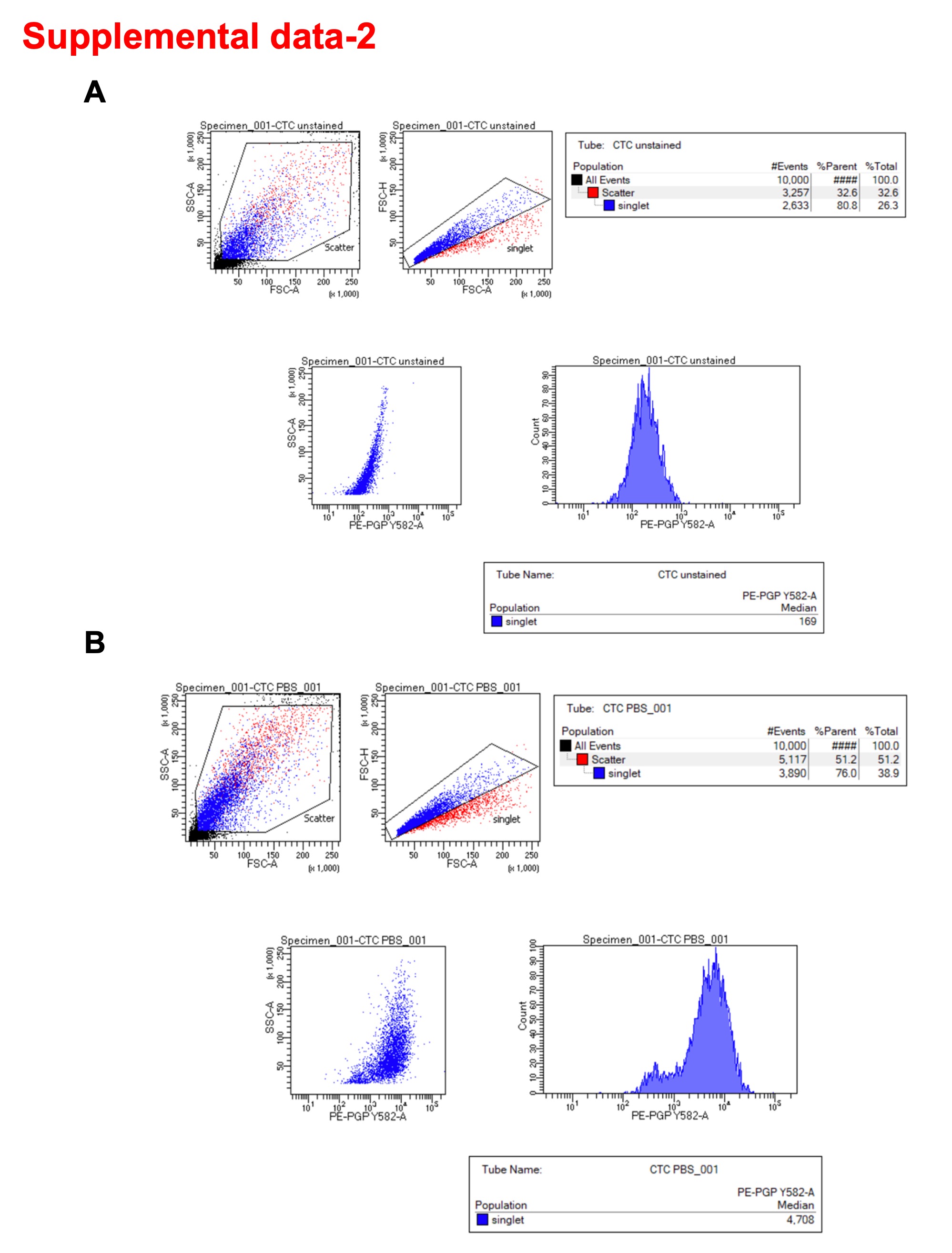

### Supplementary figure 3

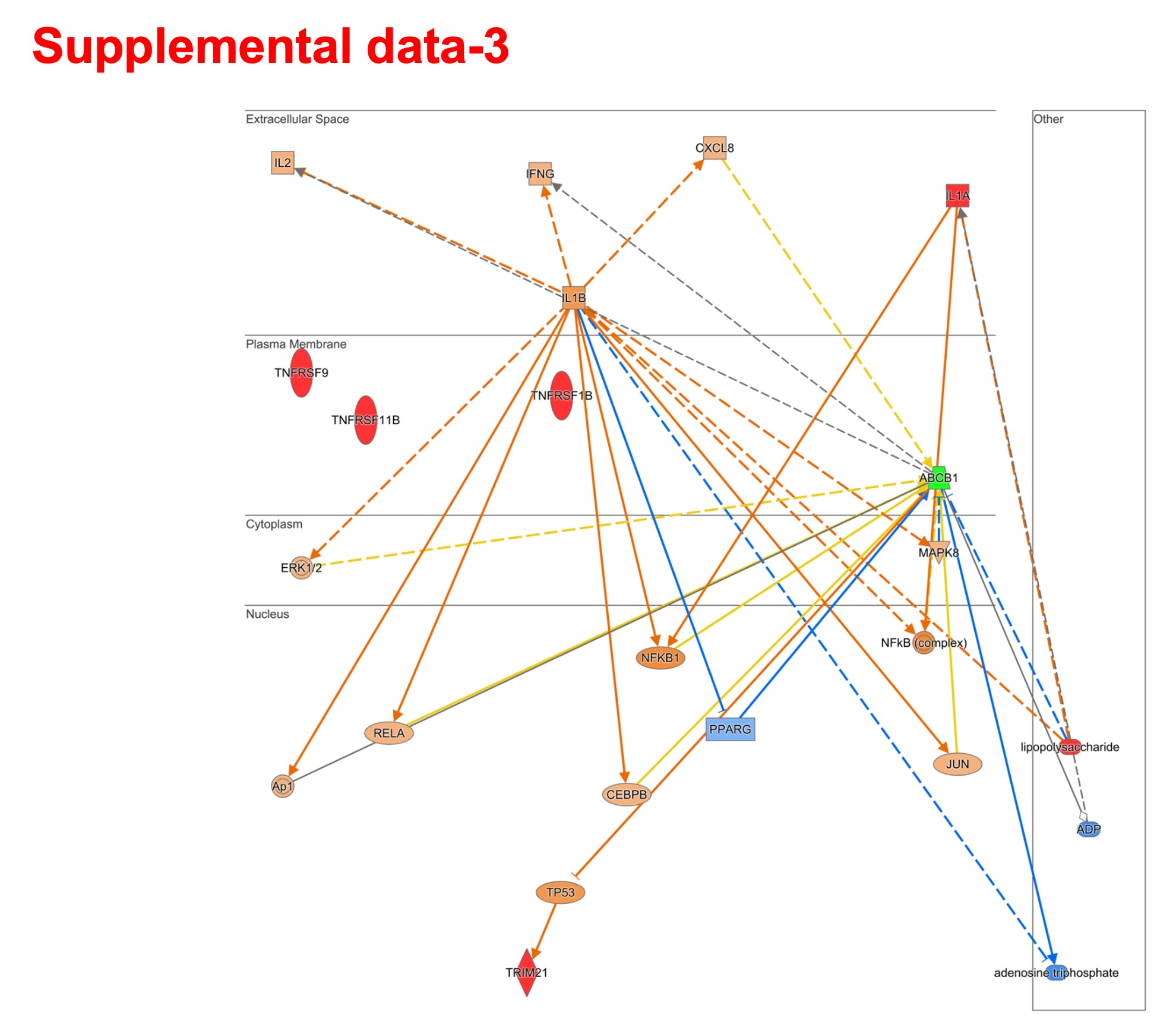

### Supplementary figure 4

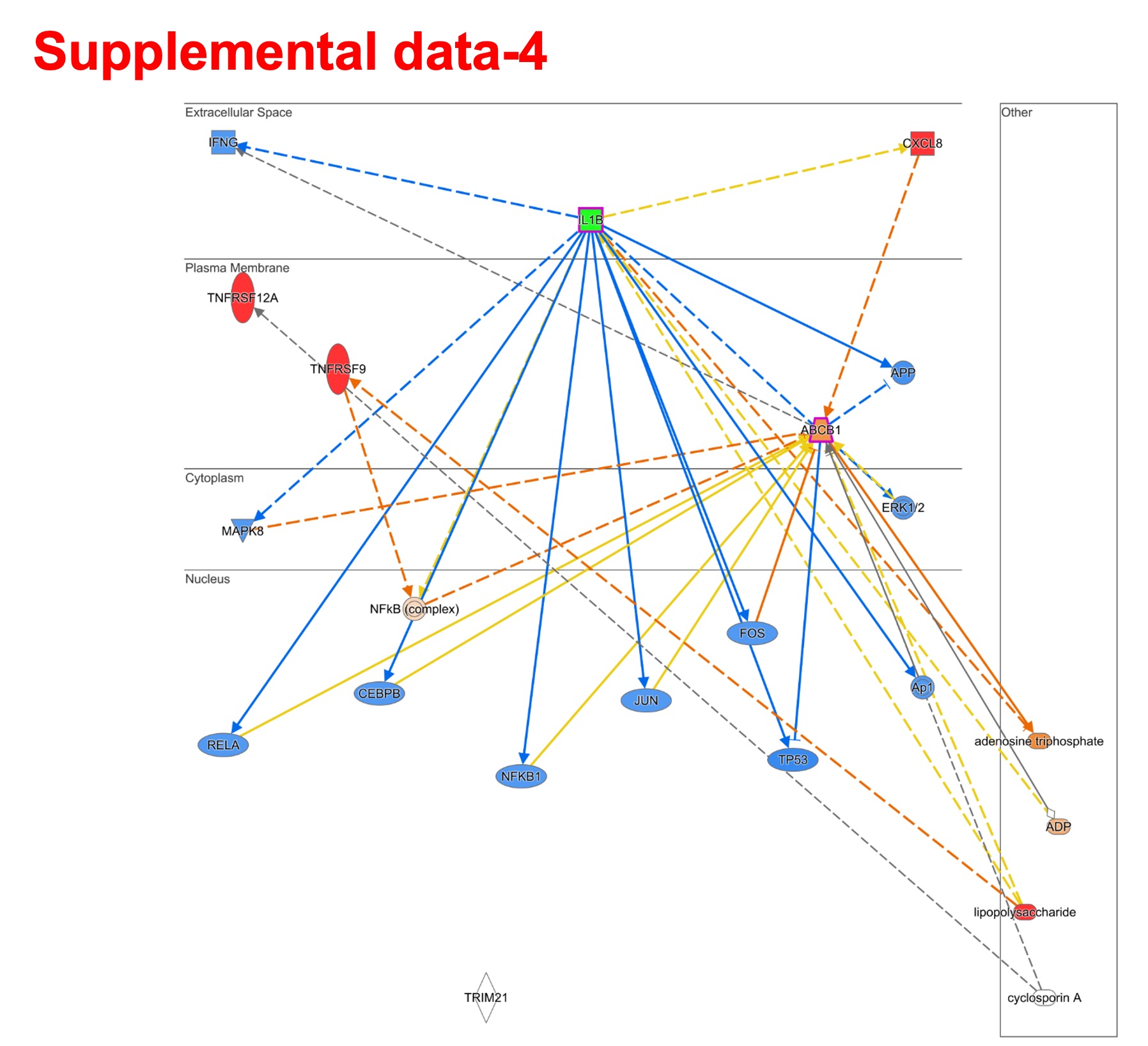

### Supplementary figure 5

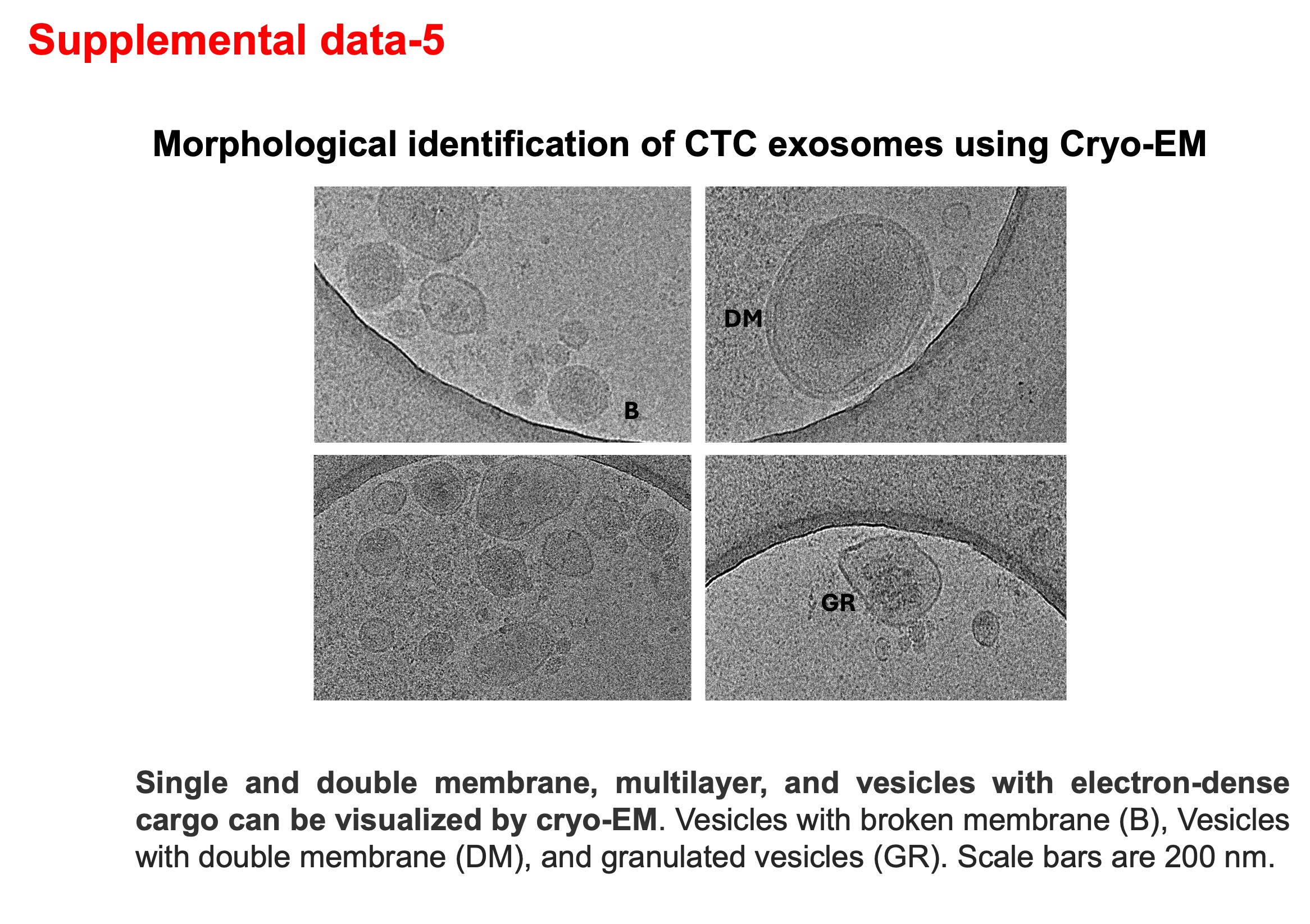

### Supplementary figure 6

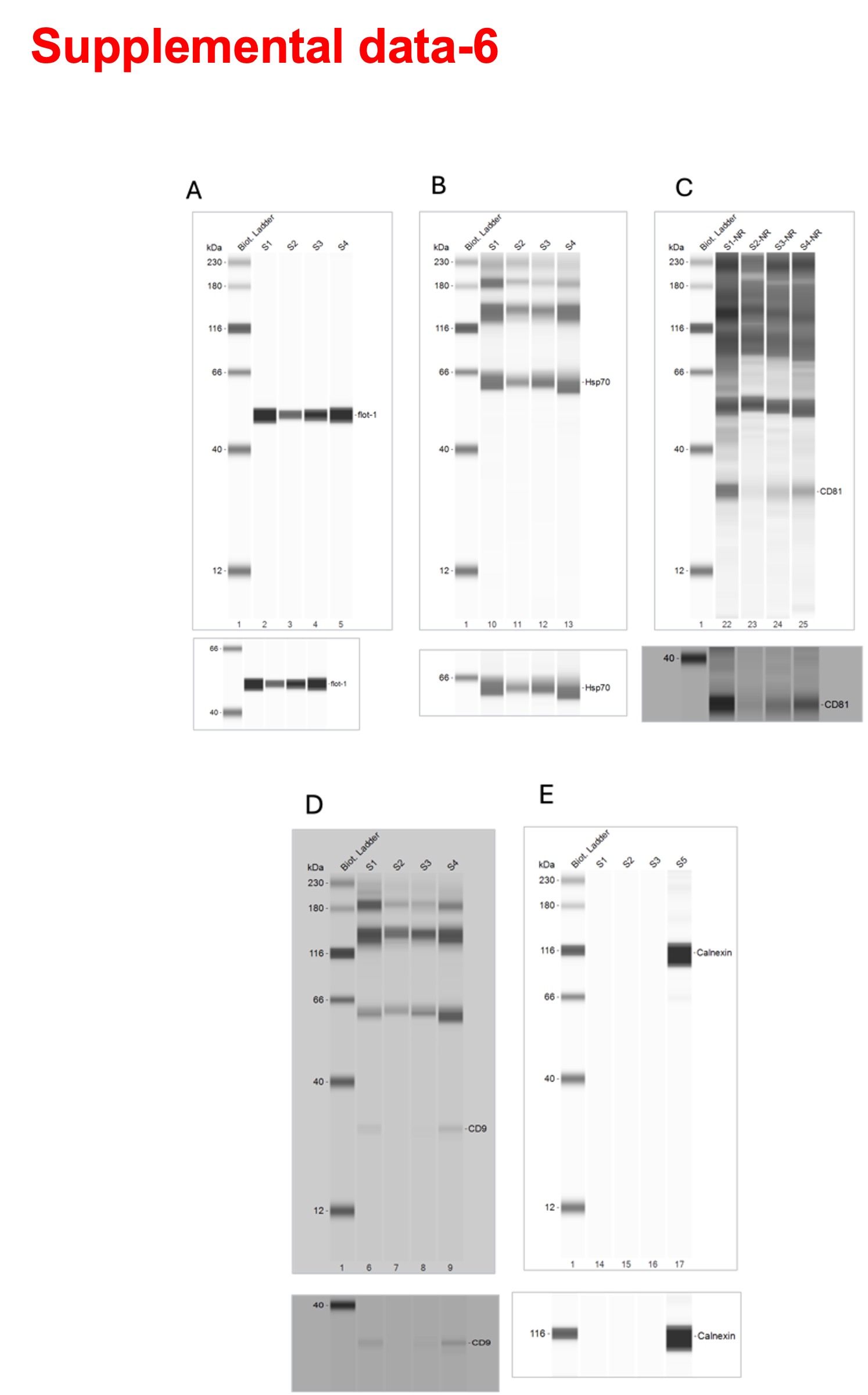

### Supplementary figure 7

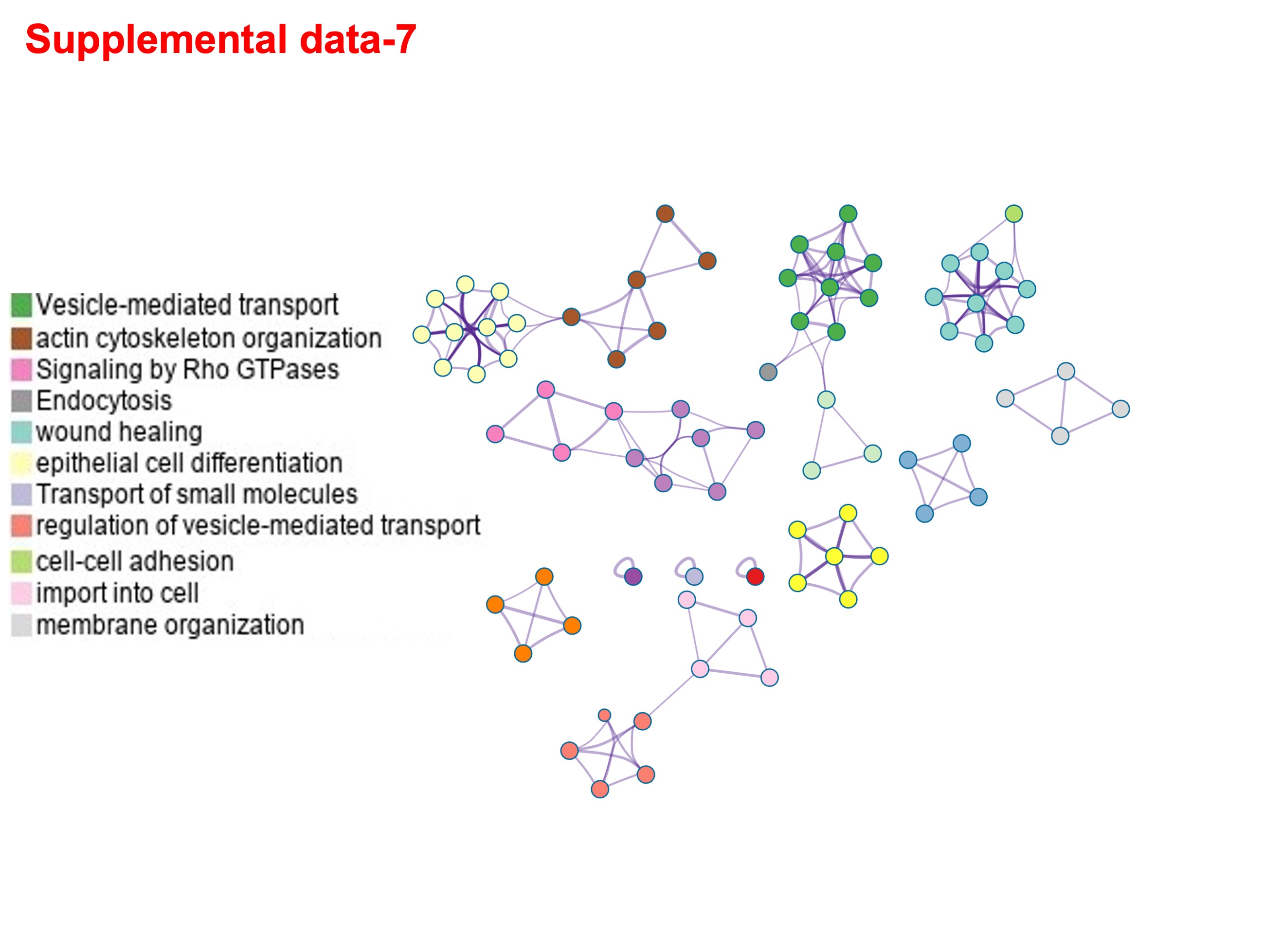

### Supplementary figure 8

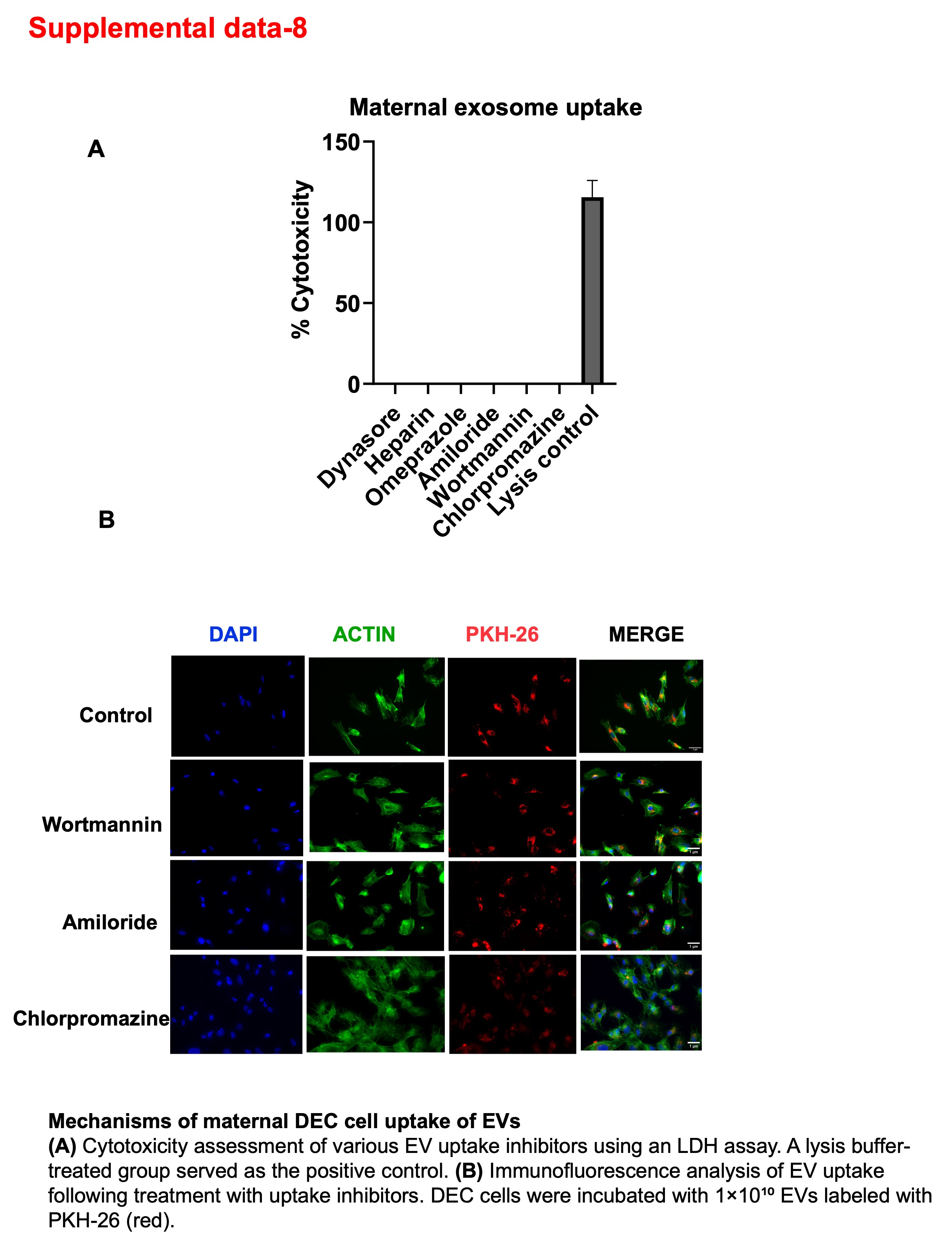

### Supplementary figure 9

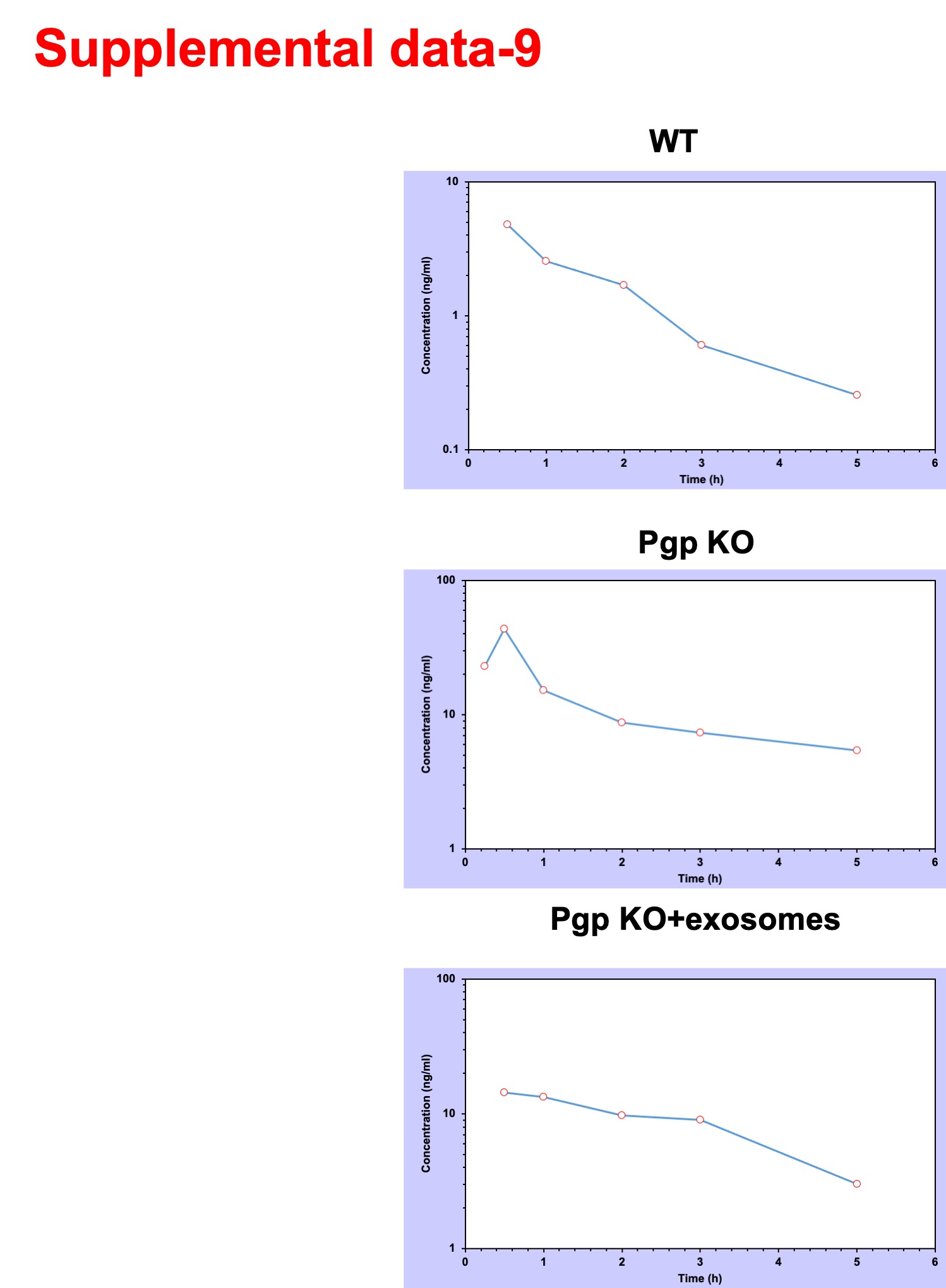
