## Supplementary Figures for "P-glycoprotein exofection between fetal and maternal cells as a mechanism of intercellular material transfer at the feto maternal interface"

**Supplementary figure _1 : Flow cytometry gating strategy for DEC with unstained control for P-gp-PE**

A. Indicates the unstained control for PE. The first gate shows a Forward scatter area (FSC-A) vs. side scatter area (SSC-A) plot for the main/total cell population (Scatter). The second gate is FSC-A vs. forward scatter height (FSC-H) for singlet populations and for removing aggregates or double cells. The table summarizes the total events for singlets, and % the total population. A histogram represents P-gp fluorescence intensity and calculates the mean fluorescence intensity. B. Represents the same gating for P-gp-stained cells, showing a positive shift towards right.

**Supplementary figure _2 : Flow cytometry gating strategy for CTC with unstained control for P-gp-PE**

A. Indicates the unstained control for PE. The first gate shows an FSC-A vs. SSC-A plot for the main cell population (Scatter). The second gate is FSC-A vs. FSC-H for singlet populations and for removing aggregates and double cells. The table summarizes the % total events were singlets, from total population. A histogram represents P-gp fluorescence intensity and calculates the mean fluorescence intensity. B. Represents the same gating for P-gp-stained cells showing a positive shift towards right.

**Supplementary figure _3: IPA analysis of the subcellular localization of gene networks associated with ABCB1 downregulation in DECs:** The IPA-derived network analysis shows upstream regulators, transcription factors, cytokines, and signaling molecules involved in the *ABCB1* downregulation in DEC in LPS treatment. The node color indicates gene expression: Green- downregulated, red- upregulated, white- unchanged. The colored lines indicate orange activation, blue inhibition, yellow lines inconsistent with the downstream node, and gray lines with no predicted direction. Solid arrows-direct interactions, dashed arrows-indirect interactions. Subcellular localization is annotated (extracellular, plasma membrane, cytoplasm, nucleus).

**Supplementary figure _4: IPA analysis of the subcellular localization of gene networks associated with ABCB1 upregulation in CTCs**: The IPA-derived network analysis shows upstream regulators, transcription factors, cytokines, and signaling molecules involved in the *ABCB1* upregulation in CTCs in LPS treatment at Subcellular localization is annotated (extracellular, plasma membrane, cytoplasm, nucleus). The node color indicates gene expression: Green- downregulated, red- upregulated, white- unchanged. The colored lines indicate orange activation, blue inhibition, yellow lines inconsistent with the downstream node, and gray lines with no predicted direction. Solid arrows-direct interactions, dashed arrows-indirect interactions.

**Supplementary figure _5: Single and double membrane, multilayer, and vesicles with electron-dense cargo can be visualized by cryo-EM**. Vesicles with broken membrane (B), Vesicles with double membrane (DM), and granulated vesicles (GR). Scale bars are 200 nm.

**Supplementary figure _6: Exosomes characterization by simple Western Jess (automated western system) A to D:** S1-S4, exosome samples were tested for the canonical exosome markers Flotillin-1, HSP70, and the tetraspanin markers CD81 and CD9 (S1NR-S4NR: exosome samples prepared in non-reduced conditions optimized for CD81 marker blot **C**). **E**: CTC Cell lysate was used as a positive control for Calnexin, an exosome purity marker, along with exosome samples.

**Supplementary figure _7: Gene ontology enrichment analysis generated by Metascape:** The indicated nodes involved in different biological processes are represented with color codes on the left. The network analysis shows dense interconnection between cellular transportation, cytoskeleton remodeling, and membrane dynamics. These CTC exosomes, which are involved in paracrine signaling and barrier function at FMi, show cell adhesion, epithelial differentiation, vesicle-mediated signaling, endocytosis, and cell import pathways.

**Supplementary figure _8: Mechanisms of maternal DEC cell uptake of EVs**
**(A)** Cytotoxicity assessment of various EV uptake inhibitors using an LDH assay. A lysis buffer-treated group served as the positive control. **(B)** Immunofluorescence analysis of EV uptake following treatment with uptake inhibitors. DEC cells were incubated with 1×10¹⁰ EVs labeled with PKH-26 (red).

**Supplementary figure _9: Drug elimination efficiency of CTC exosomes with P-gp as a cargo**. Semi-logarithmic plot of serum drug concentration on the y- axis versus time indicated on the x- axis. WT showing one-compartment kinetics and a smooth drug elimination (monoexponential decline). P-gp KO Shows the initial sharp decline in the drug concentration, followed by a slower elimination, indicating drug distribution and slow elimination following the two-compartmental drug kinetics (Biphasic decline). P-gp KO+ exosomes Gradual decline shows steady mono-exponential drug elimination with one-compartment kinetics as in P-gp WT mice (monoexponential decline).
